# Time- and cell-dependent atypia and cell death are caused by progressive deficiency in DNA replication

**DOI:** 10.1101/2020.12.31.425001

**Authors:** Alex Y. Lin, Georgia K. Thomas, Khai Chung Ang, Damian B. van Rossum, Victor A. Canfield, Keith C. Cheng

## Abstract

Pleiotropy caused by single-gene mutations is common and poorly understood. A zebrafish null mutant of DNA polymerase α subunit B, *huli hutu* (*hht)*, evolves a complex pleiotropy associated with DNA damage and S phase arrest across multiple organ systems over 5-7 days, including nuclear atypia, a common cellular feature in human cancers and pre-cancers, in gastrointestinal organs, and nuclear fragmentation in the eye and brain. The pleiotropic pattern of *hht* phenotypes is explained by progressive loss of wild-type maternal *pola2* function in homozygous mutant embryos whose *pola2* mRNA becomes undetectable by 24 hours post-fertilization (hpf). Inhibition of DNA synthesis by aphidicolin or hydroxyurea in wild-type embryos from 24 hpf phenocopied the pleiotropic pattern of *hht*. These results are consistent with a model in which time-sensitive, reduced capacity for DNA synthesis results in cell death in fast-replicating cells, and nuclear atypia in tissues with fewer and larger cells.

## Introduction

Changes in nuclear morphology occur during normal cellular processes such as mitosis, differentiation, and cell migration (Jevtić *et al*. 2014, Calero-Cuenca *et al*. 2018), but histologically-defined cellular atypia, characterized by darker hematoxylin staining and large, irregular nuclear shape, is common in both neoplasia and in settings that involve physical or chemical damage to DNA, such as UV-irradiation and chemotherapy (Blum 1978, Stenbäck 1978, Carr and LiVolsi 1989, Kim *et al*. 2016). Nuclear atypia is also used as a diagnostic signature in many human cancers and pre-cancers (Rosai 2004, Billings and Goldblum 2010, Kumar *et al*. 2010, Lanzkowsky *et al*. 2016, Pizzorno *et al*. 2016). The presence and severity of nuclear atypia generally correlates with higher tumor grade and poor prognosis (Rosai 2004, Kodota *et al*. 2014, Manimaran *et al*. 2014, Yamaguchi *et al*. 2015, Poropatich *et al*. 2016, Zhou 2018). Despite the clinical diagnostic relevance of nuclear atypia, its genetic and mechanistic origins are poorly understood.

The small size of zebrafish larvae allowed us to generate histological arrays for pursuit of a histology-based forward genetic screen to identify genes that can cause nuclear atypia (Mohideen et al. 2003). Our laboratory identified a recessive larval-lethal mutant, *huli hutu* (*hht*), that develops optical opacity of the brain, small eyes, an upward curvature of the body, and dies after 5 to 7 days of development. Histology and X-ray histotomography revealed that virtually all organs and tissues exhibited severe cytological abnormalities (Mohideen *et al*. 2003, Ding *et al*. 2019). Highly proliferative tissues, particularly of the retina and brain, and to a lesser extent, the gastrointestinal tract organs, were affected most dramatically.

Positional cloning shown here revealed a frameshift mutation in *pola2* in *hht*, which encodes the B subunit of the DNA polymerase α-primase complex (Pol α). Immediate growth arrest phenotypes associated with *pola2* mutation in two other organisms, *Saccharomyces cerevisiae*, (Foiani *et al* 1994) and *Arabidopsis thaliana* (Yang *et al*. 2009) begged the question of the reason underlying the longer survival of zebrafish *pola2* mutants. In *Caenorhabditis elegans*, mutation of the *pola2* paralog *div-1* substantially increased the duration of interphase and produced embryos that failed to hatch or to generate intestinal or pharyngeal cells (Encalada *et al*. 2000). We detected wild-type maternal *pola2* mRNA in homozygous mutant embryos, providing an explanation for the extended lifespan of *pola2* mutant zebrafish. Consistent with the disappearance of detectable *pola2* mRNA at 24 hpf, two independent chemical means of inhibiting DNA synthesis beginning at 24 hpf phenocopied cell-specific *hht* phenotypes. Here, we show correlations between replicative demand and DNA damage that appear to explain the disparity in mutant phenotype between cell types. This work also sheds light on potential mechanisms of pleiotropy, which are being increasingly recognized as common in biology and disease (Goh *et al*. 2007, Ittisoponpisan *et al*. 2017).

## Results

### A zebrafish mutant with abnormal nuclear morphology

The gross phenotype of *hht* mutants is evident at 3 days post-fertilization (dpf) under a dissecting microscope and includes reduced head and eye size, a curved body, and a pronounced yolk (Mohideen et al. 2003, **Fig. 1A**). This gross phenotype persists until death, which occurs between 5 and 7 dpf (**Fig. 1B**). A detailed examination by histology at 5 dpf showed that while all organs and tissues were distinguishable, virtually all exhibited severe cytological changes (**Fig. 1C**). Highly proliferative tissues, including retina, brain, and gastrointestinal tract organs, were affected most dramatically. Enterocytes in the *hht* intestinal lumen were irregular in polarity, size and shape, cellular and nuclear boundaries were difficult to define, and cell nuclei were often hyperchromatic and contained prominent nucleoli. Other endodermally-derived organs such as pancreas and liver also exhibited severe cytologic atypia, often presenting with enlarged nuclei and high variability in cellular and nuclear size and shape. Dysmorphologies were also detected in cartilage, where the normal aligned arrangement of cartilage cells in neat rows (in wild-type siblings) is replaced by cells with variable shapes, sizes, and polarity, and irregular arrangements. The eyes of *hht* mutants have misshapen lenses and retinas with a substantial reduction in cell number, a loss of stratification of retinal layers, and nuclear fragmentation. Further characterization using the interactive x-ray histotomography zebrafish database, 3d.fish, revealed stark contrast in phenotype between dividing and nondividing cells: dying cells in the retina vs. well-aligned, differentiated sensory epithelium (**Fig. 2**, Ding *et al*. 2019, Dyballa *et al*. 2017).

**Figure 1.**
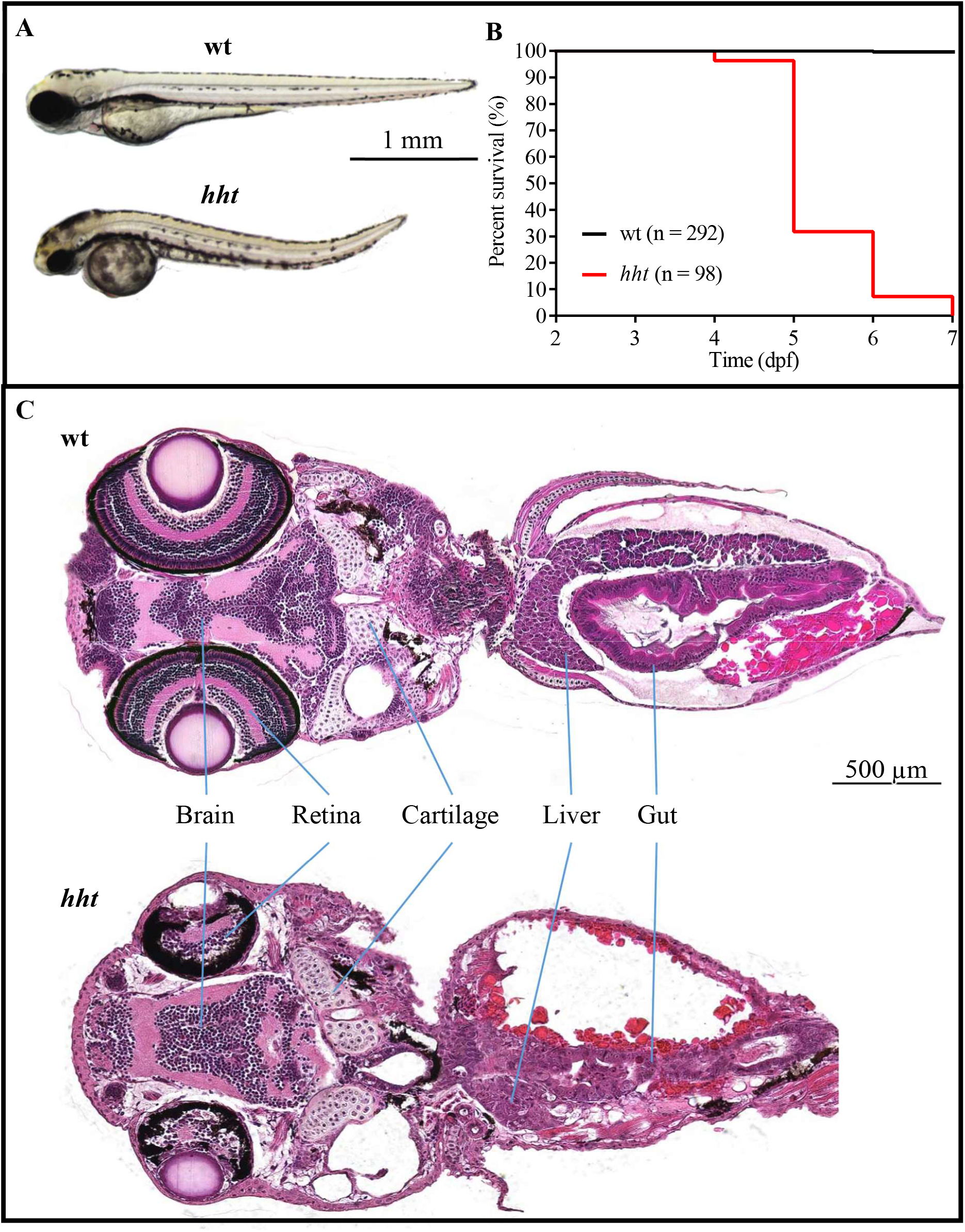
The pleiotropic *hht* mutant survives up to 7 days. **A)** The *hht* mutant gross phenotype includes reduced head and eye size, an enlarged yolk, and dorsal curvature of the tail. **B)** The *hht* mutants show increasing larval lethality over 7 days; none survive past 7 days. In the same time period, only 1 wild-type sibling died. **C)** 5 dpf *hht* mutants show disruption in tissue organization and a range of cellular dysmorphologies across most cell types.

**Figure 2.**
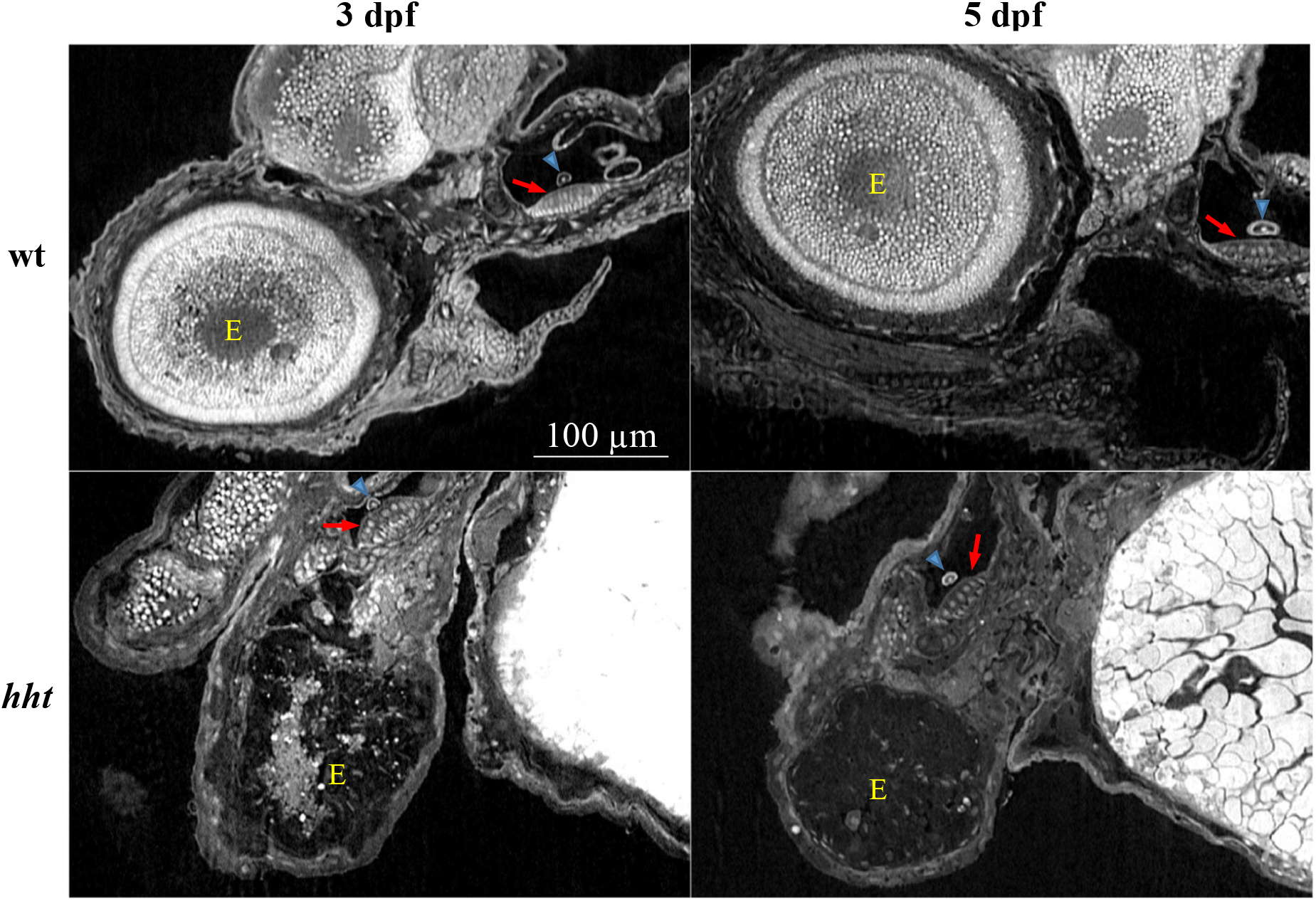
X-ray histotomography of eye and sensory epithelium in wild-type and *hht* larvae. Images of the eye and sensory epithelium of wild-type and *hht* fish at 3 and 5 dpf are extracted from the interactive 3d.fish database developed by Ding *et al*. (links in Materials and Methods). Retinal cells, which replicate quickly to large numbers in wild-type larvae, exhibit drastic cell loss accompanied with nuclear fragmentation in *hht* mutants. The sensory epithelium, which becomes fully differentiated and consists of a small number of cells, appears normal and organized in both wild-type and mutant larvae. Red arrows = sensory epithelium; blue arrow = otolith. Scale bar = 100 µm.

### Pola2 mutation is responsible for the hht phenotype

#### Positional cloning of hht

Microsatellite-based positional cloning and gene knockdown by morpholino oligonucleotides was used to show that the causative mutation for the *hht* phenotype is in *pola2*. The identification of zero recombinants in 1948 meioses with markers *z10868* and *z15236* revealed that the gene responsible for the *hht* phenotype was within 0.05 centimorgans (cM) of these markers, which corresponds to about 35.5 kb in zebrafish (Shimoda *et al*. 1999). These two markers were flanked on either side by microsatellite markers *z26580* and *z13225*, each of which had one recombinant in 1948 meioses, indicating that the causative mutation resides between these markers (**Fig. 3A**). Five candidate genes, *cenph, dimt1l, mier3b, mrps36*, and *pola2*, are annotated in this region.

**Figure 3.**
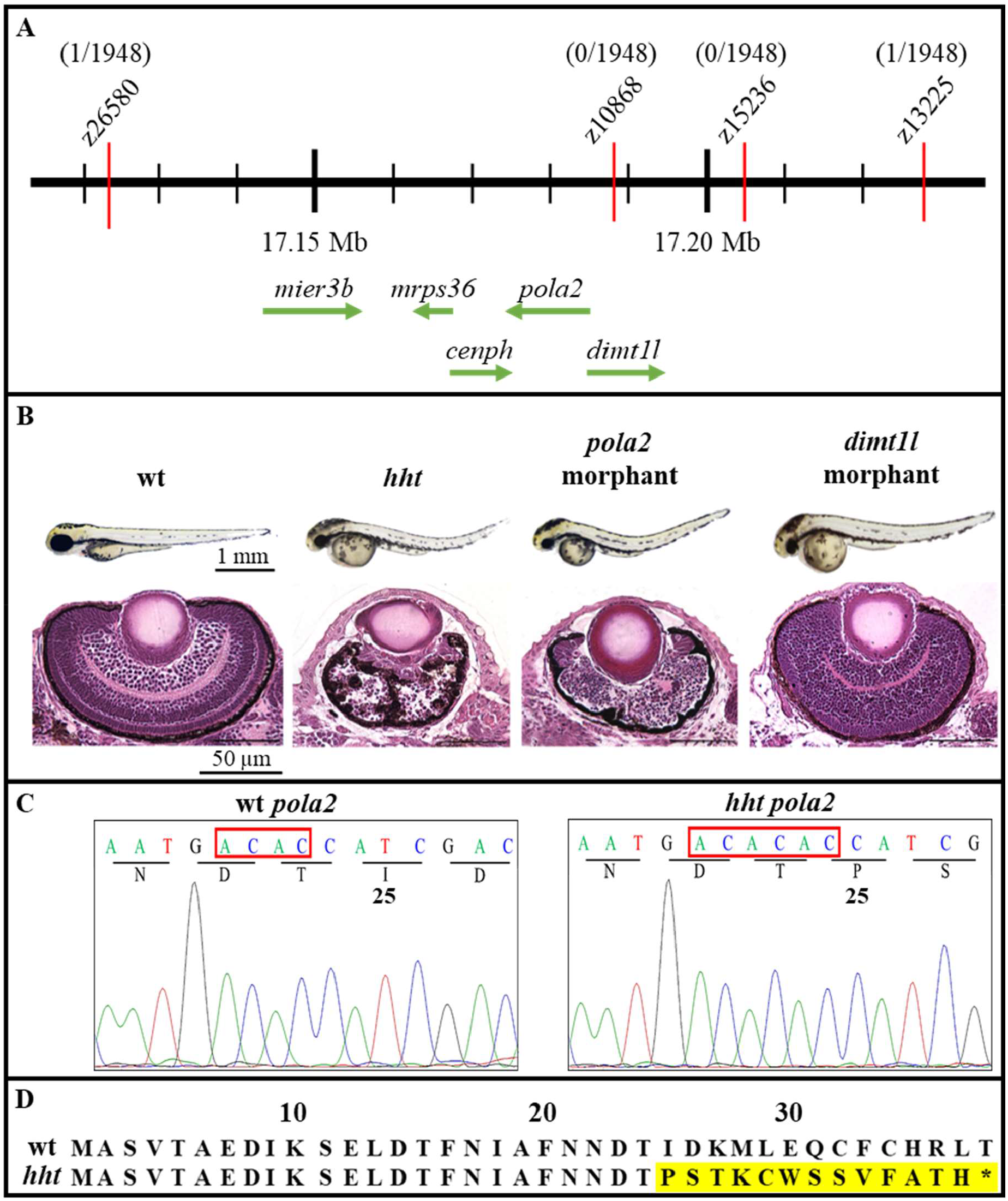
Microsatellite-based positional cloning of *pola2*. **A)** In positional cloning, the frequency of co-segregation of a phenotype of interest and polymorphic markers is used to locate the causative mutation. The map distance between the causative mutation and the markers is calculated from the frequency of recombinants as indicated above each marker. For markers *z26580* and *z13225*, 1 recombinant out of 1948 meiosis corresponds to ∼ 0.05 centimorgans (cM), or 32.5 kb, in zebrafish. The lack of recombinants for markers *z10868* and *z15236* indicates that the causative mutation is physically closer to these two markers and resides in the interval between *z26580* and *z13225*. Five candidate genes, *mier3b* (mesoderm induction early response factor 1, family 3 b), *mrps36* (28S mitochondrial ribosomal protein S36), *cenph* (centromere protein H), *pola2* (DNA polymerase alpha, subunit B), and *dimtl1* (dimethyladenosine transferase), were annotated in the zebrafish genomic database in the identified GRCz11 region. **B)** Gross and histological examination of the eyes of wild-type, *hht, pola2* morphant, and *dimt1l* morphant revealed a striking similarity in reduced cell number, loss of stratification, thickened corneal epithelium, and disruption in the pattern of retinal pigmented epithelial pigment between *hht* and *pola2* morphant, suggesting that *pola2* was the affected gene in *hht*. **C)** a 2-nucleotitde AC insertion in the double AC repeat starting at the 68^th^ nucleotide was detected in *hht*. **D)** This frameshift mutation results in changes in amino acid sequence starting at the 25^th^ amino acid, ending with a premature stop codon at the 38^th^ amino acid position. (red box = AC insertion; yellow highlight = amino acid change; asterisk = stop codon).

To identify the gene responsible for the *hht* phenotype, one translational blocking and two splice-junction blocking morpholino oligonucleotides (MOs) were designed per candidate gene and injected into wild-type zebrafish embryos at the one-cell stage to inhibit translation or RNA splicing, respectively. Only *pola2* and *dimt1l* morphants exhibited a combination of reduced head and eye size, a pronounced yolk, and a dorsal curvature that were grossly similar to *hht* (**Supplemental S1**). Histological examination of cellular phenotype at 3 dpf revealed striking similarities between *pola2* morphants and *hht*, both demonstrating reduced retinal volume and cell number, loss of retinal layers, fragmented cell nuclei, and a thickened uneven corneal epithelium (**Fig. 3B**). The eyes of *dimt1l* morphants, while distinctly abnormal compared with wild-type zebrafish, retained the overall size, ovoid shape, some retinal layering, and cell types of wild-type eyes, and lacked the profound deficiencies found in *hht* eyes. Taken together, mapping data and the near identity in phenotypes of *hht* and *pola2* morphants indicate that the affected gene in *hht* is *pola2*.

Sequencing of gDNA and cDNA of wild-type and mutant *pola2* revealed a 2-nucleotide AC insertion at the double AC repeat at the 68^th^ nucleotide from the translation start site (**Fig. 3C**). The frameshift mutation results in a premature stop codon at the 38^th^ amino acid position (**Fig. 3D**), suggesting that the final protein product is truncated and non-functional. The AC insertion, located at the 2^nd^ exon of the 19-exon zebrafish *pola2* gene, is therefore most likely the causative mutation. To verify this insertion, genomic DNA from 20 each of *hht* larvae and wild-type larvae from wild-type siblings of their parents was pooled to sequence PCR amplicons of the 2^nd^ exon of *pola2;* the 2-nucleotide insertion was present in the *hht* pool, but not the wild-type pool.

#### Partial rescue of the huli hutu mutant phenotype with wild-type pola2 mRNA

One prediction of *pola2* being the affected gene in *hht*, is that injection of wild-type mRNA would rescue at least some of the several phenotypes of *hht* fish. Wild-type *pola2* mRNA was transcribed from a cDNA clone from the wild-type Connors background strain and injected into embryos from 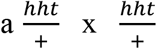 cross at the 1-cell stage with the expectation of a 1:2:1 ratio of 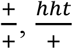, and 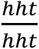 genotypes (**Supplemental S2**). At 3 dpf, 90% of injected 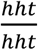 larvae, identified by genotyping, were partially rescued, exhibiting an intermediate gross phenotype – a straight body and yolk resembling that of a wild-type, but with eyes and brain that are smaller than wild-type but larger than *hht* (**Table 1, Fig. 4**). This intermediate phenotype has not been seen in hundreds of crosses between *hht* heterozygotes. All larvae with an intermediate phenotype were genotypically homozygous mutant 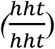. Retinal cell layering of the rescued fish was reminiscent of those seen in homozygous wild-type larvae. The *hht* intestinal epithelial phenotype, in which cell nuclei show loss of polarity and prominent nucleoli, was absent in mRNA-rescued *hht* larvae. Fragmented nuclei in the brain and eyes of *hht* larvae were also absent in both wild-type and rescued fish. The incomplete rescue phenotype can be explained by an expected degradation and/or dilution of the injected wild-type *pola2* mRNA over time, leading to a slower rate of cell division than in wild-type fish.

**Table 1:** Wild-type *pola2* mRNA rescues *hht* mutants

| Injected mRNA | Total | $\frac{+}{+}$ , $\frac{hht}{+}$ | $\frac{hht}{hht}$ | |
| --- | --- | --- | --- | --- |
| | | Phenotypically wild-type / (expected) | Partially rescued / total $\frac{hht}{hht}$ | Phenotypically *hht* |
| none | 182 | 136 / (137) | 0 / 46 | 46 |
| wt *pola2* RNA | 302 | 234 / (227) | 61 / 68 | 7 |
| mutant *pola2* RNA | 75 | 60 / (56) | 0 / 15 | 15 |

**Figure 4.**
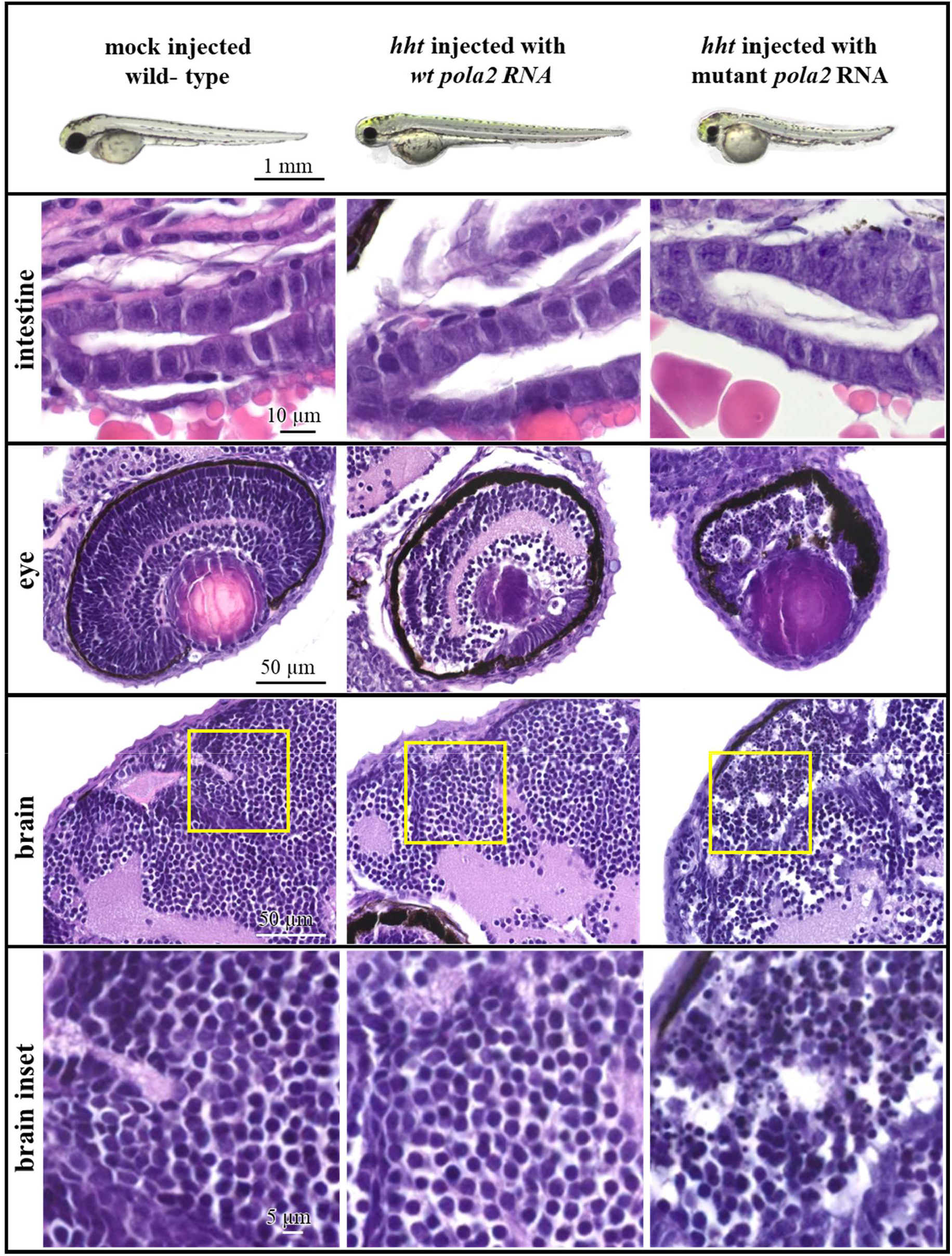
Wild-type mRNA partially rescues the *hht* phenotype. Rescued *hht* mutants exhibit a straight body and a normal yolk. Sizes of the eyes and head are intermediate between wild-type and *hht* mutants. Rescued *hht* fish show histologically normal intestinal epithelium and partial preservation of the retinal layering found in wild-type eyes. The eyes and brain of partially rescued *hht* fish injected with wt mRNA show no evidence of nuclear fragmentation. Histology sections were imaged at 63X.

A mixture of wild-type and mutant embryos from a 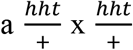 cross were injected at the 1-cell stage with phenol red, 200 pg wild-type *pola2* mRNA, or 200 pg mutant *pola2* mRNA. Number of embryos injected with no mRNA or wild-type *pola2* mRNA was aggregated from three separate experiments.

Another prediction of *pola2* mutation being responsible for the *hht* phenotype is that knockout of *pola2* on both chromosomes by genome editing would show a *hht* phenotype. Wild-type larvae were rendered mutant for *pola2* using the CRISPR/Cas9 genome editing system. Homozygous knockout fish harboring a 16-nucleotide insertion between nucleotides 927-928 (**Supplemental S3A**) exhibited a gross phenotype that was strikingly similar to *hht* (**Supplemental S3B**). The frameshift in this CRISPR insertion allele disrupts the protein sequence after amino acid 308. Under the assumption that human and zebrafish POLA2 proteins share similar structures, these *pola2* knockout zebrafish would produce a mutant protein without a functional phosphodiesterase (PDE) domain, which has been reported to be important for binding between human POLA2 and the catalytic A subunit of Pol α, predicting a null phenotype (Klinge *et al*. 2009, Suwa *et al*. 2015). The gross phenotypes of *hht* and *pola2* knockout larvae were indistinguishable, further confirming that the affected gene in *hht* is *pola2*.

We have established that *hht* is mutant in the *pola2* gene by positional cloning, morpholino knockdown to generate a similar histological phenotype, CRISPR knockout, and partial rescue with *pola2* mRNA. Human POLA2 protein, the B subunit of DNA polymerase alpha, is known to mediate binding of the A catalytic subunit and the chromosome (Suwa *et al*. 2015). As predicted by this model, insertional mutant of the A subunit, encoded by *pola1*, resulted in gross and histological brain and eye phenotypes indistinguishable from *hht* mutants (**Supplemental S4**).

### pola2-deficient zebrafish mutants exhibit reduced DNA synthesis

Since *pola2* plays a critical role in DNA replication, we investigated how DNA synthesis was affected in the *hht* mutants. Wild-type and *hht* larvae between 2 to 5 dpf were assessed for DNA replication by incorporation of 5-ethynyl-2’-deoxyuridine (EdU), a synthetic thymidine analog. After 30 min of EdU incubation, wild-type larvae exhibited strongly positive EdU staining at all ages examined. In contrast, *hht* larvae were EdU-negative from 2 to 5 dpf, indicating a reduction of DNA synthesis below the limits of detection (**Fig. 5**).

**Figure 5.**
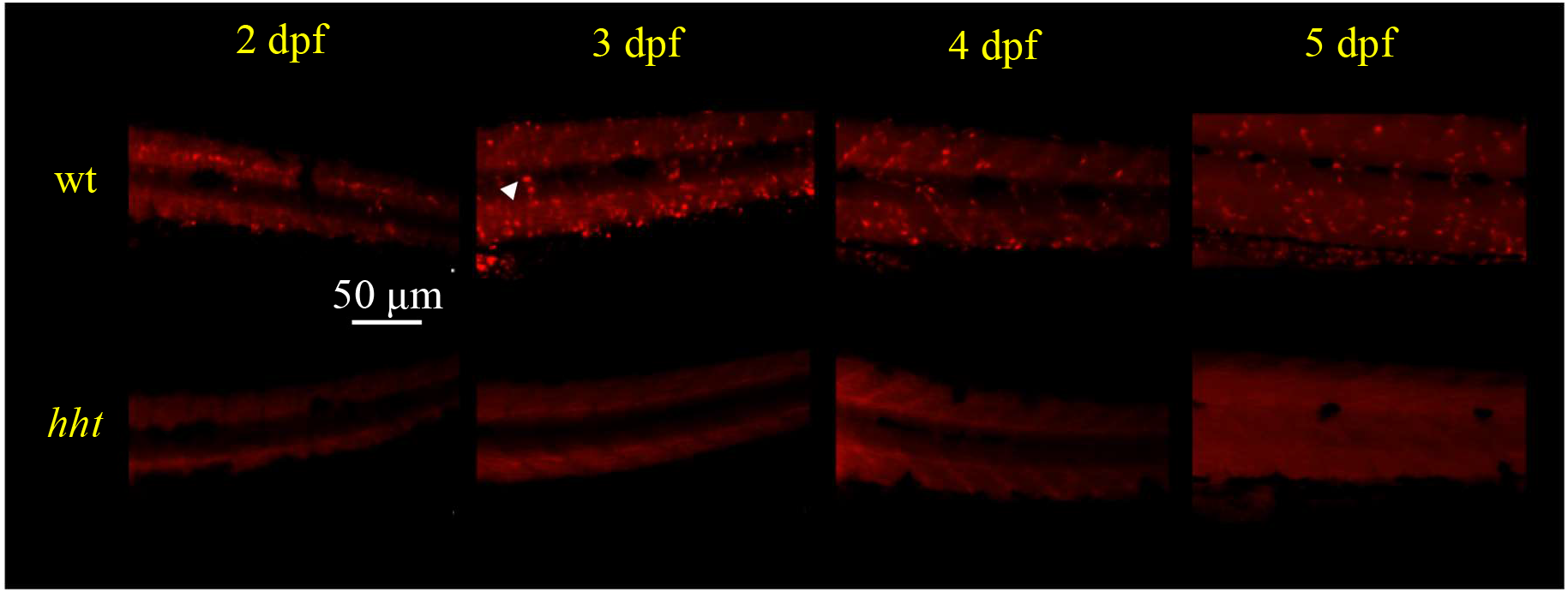
DNA synthesis is reduced in *hht* larvae. Representative images of wild-type and *hht* larvae stained with EdU. EdU-positive staining was observed in wild-type larvae, but absent in *hht* larvae at 48, 72, 96, and 120 hpf. 3 larvae were examined for each condition. White arrow = a positive focus.

The observed global reduction of DNA synthesis in *hht* mutants suggests a model in which inhibition of DNA synthesis by any mechanism can potentially reproduce the *hht* phenotype, which includes largely normal development up to 24 hpf followed by progressive dysmorphology and cell death. Based on this timing, we hypothesized that chemical inhibition of DNA synthesis by multiple mechanisms may be able to phenocopy *hht*, if started at about 24 hpf. To test this idea, wild-type larvae were exposed to hydroxyurea, an inhibitor of deoxyribonucleotide production, or aphidicolin, an inhibitor of replicative DNA polymerases α, δ, and ε (Zhang *et al*. 2008), beginning at different times. Wild-type embryos were treated with empirically determined concentrations of 50 μM aphidicolin or 150 mM hydroxyurea to induce the *hht* phenotype. Starting inhibition at 2 hpf resulted in death by 24 hpf (**Fig. 6A**). Inhibiting DNA synthesis beginning at 24 hpf (but not before) phenocopied *hht* by 72 hpf (**Fig. 6B; Supplemental S5, S6**). Cellular phenotypes in the eyes, brain, and intestine were confirmed by histology (**Fig. 7**).

**Figure 6.**
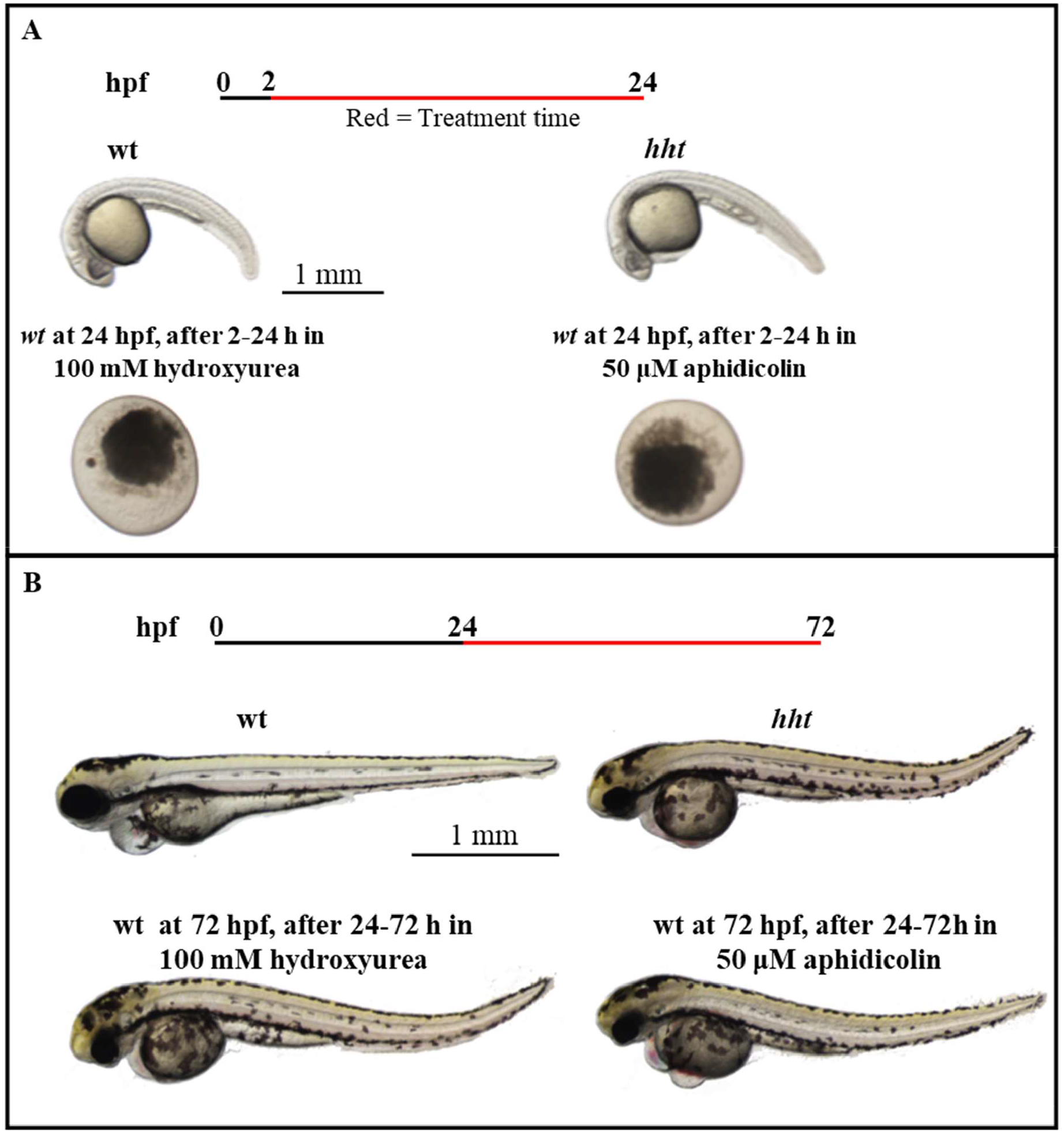
Chemical inhibition of DNA synthesis in *wt* larvae after 24 hours phenocopies *hht*. **A)** Treatment wild-type embryos with 100 mM hydroxyurea or 50 μM aphidicolin from 2 hpf on caused death by 24 hpf. **B)** Treatment of wild-type larvae (the time maternal wt *pola2* mRNA becomes undetectable) with 100 mM hydroxyurea or 50 μM starting at 24 hpf yields *hht* phenocopies at 72 hpf. Black line = no treatment; red line = treatment.

**Figure 7.**
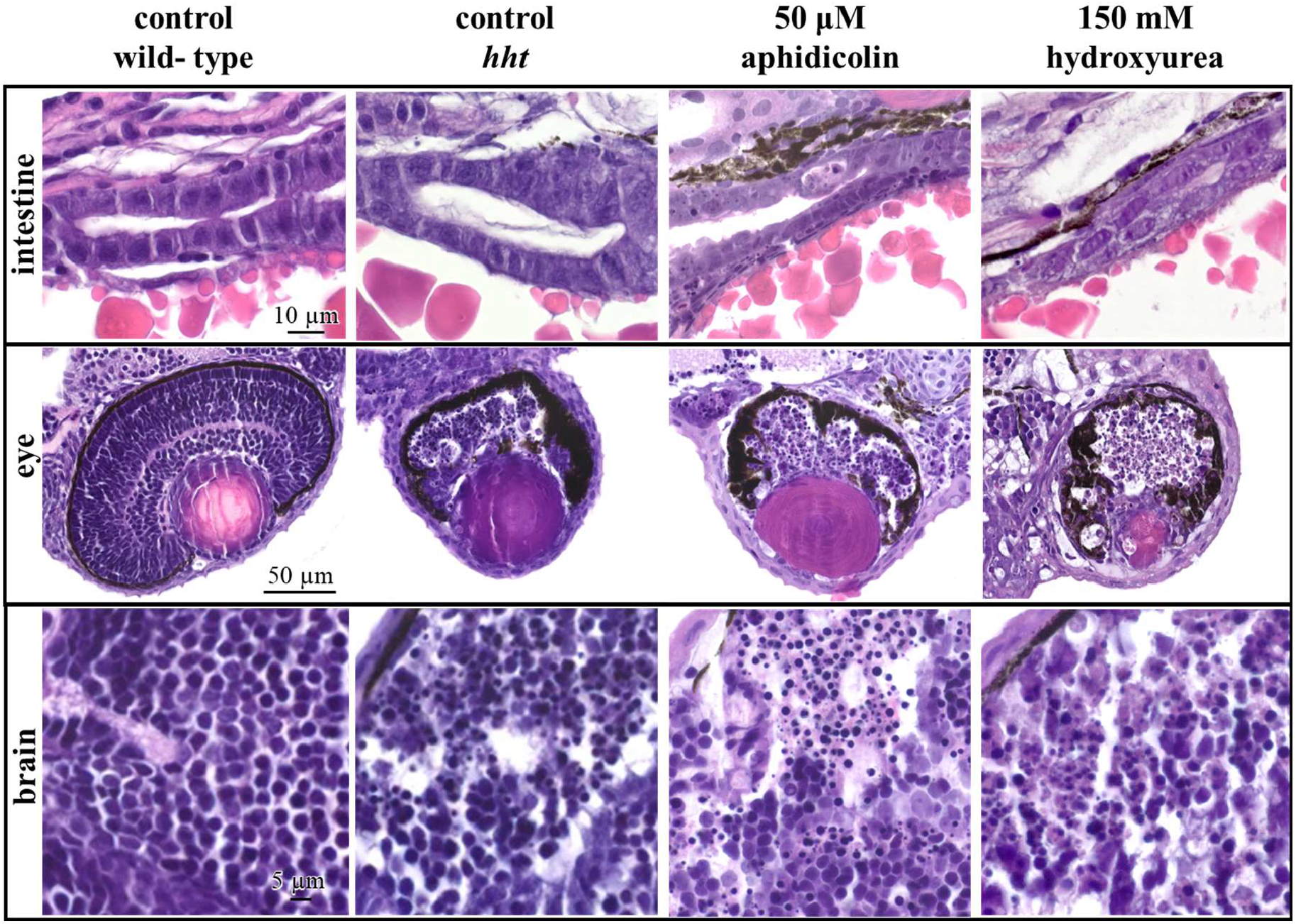
Chemical inhibition of DNA synthesis starting at 24 hpf phenocopies *hht* cellular phenotypes. Histology sections were imaged at 63X. Continuous exposure of wild-type larvae to 50 μm aphidicolin or 150 mM hydroxyurea starting at 24 hpf results in nuclear fragmentation in the prefrontal cortex and eyes. Eyes also suffer a drastic reduction in cell number and organization of retinal layers. Cellular and nuclear atypia are observed in the intestine – cell and nuclear sizes are irregular, cell boundaries are obscure, and prominent nucleoli are present.

Since inhibition of DNA synthesis predicts cell cycle arrest in S phase, higher proportions of cells are expected to be in S phase. We measured the proportion of cells in S phase by flow cytometric measurements of DNA per cell in propidium iodide-labeled cells dissociated from wild-type and *hht* larvae (**Fig. 8**). At 3 dpf, the proportion of cells in S phase was approximately 3-fold higher in *hht* (45.9 ± 5.8 %) than in wild-type (15.5 ± 3.2 %; p < 0.05, t-test). S phase accumulation in *hht* cells persisted to 5 dpf (p < 0.05, t-test). In this analysis, all cell types were examined and the proportion of cells in each phase of the cell cycle represented an average across heterogeneous cell populations.

**Figure 8.**
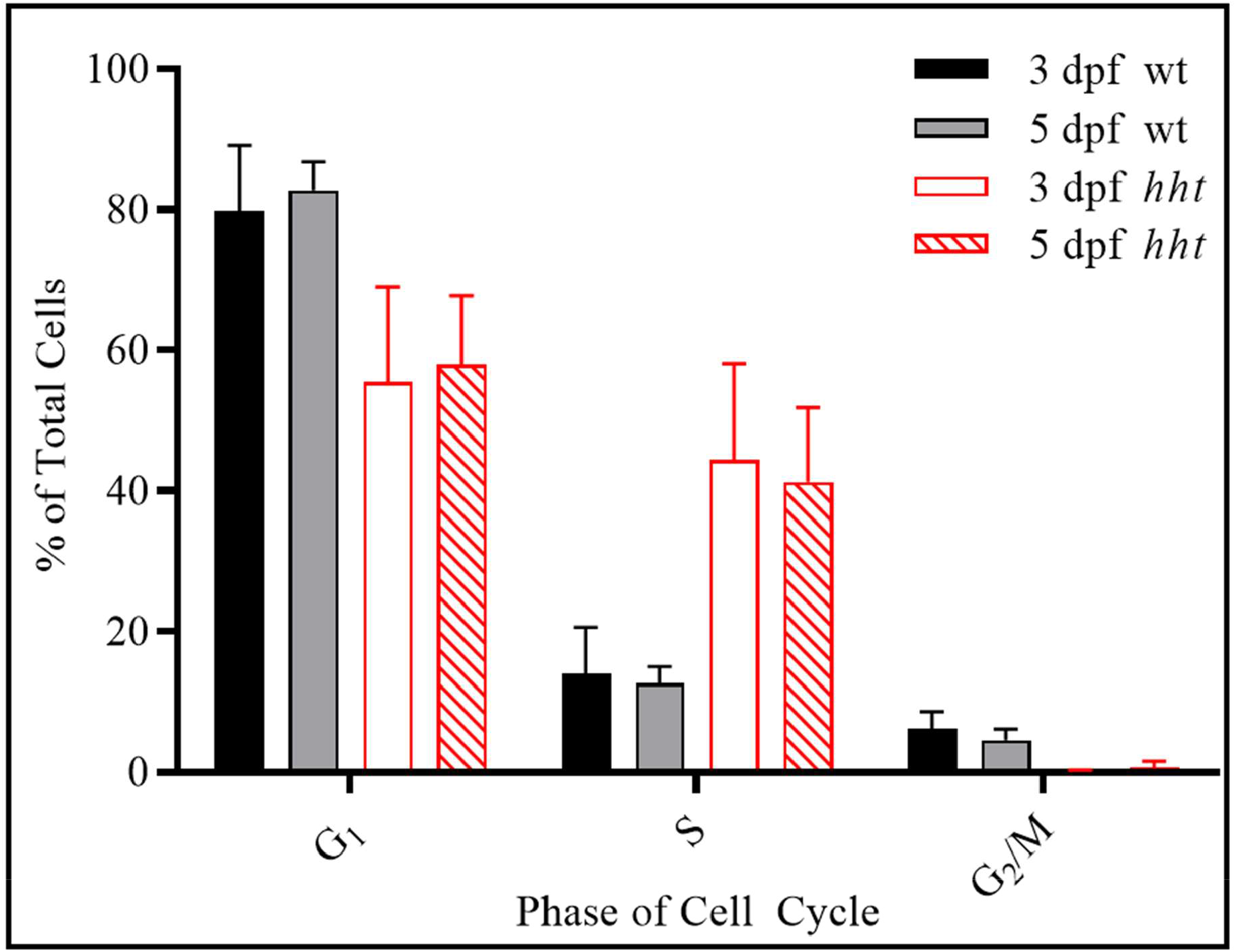
Accumulation of cells in S phase in *hht*. Wild-type and *hht* larvae were dissociated into single cells at 3 and 5 dpf, stained with propidium iodide, and analyzed by flow cytometry. Compared to wild-type, the proportion of cells in S phase increased significantly in *hht* at 3 dpf (p < 0.05) and persisted to 5 dpf (p < 0.05). Error bars = standard error of the mean.

### *Tissue specificity of cell death and DNA damage phenotypes in* hht *zebrafish*

Nuclear fragmentation as seen in the eyes of *hht* larvae indicates cell death. To assess the extent and timing of cell death, live wild-type and *hht* larvae at 24, 36, 48, 72, 96, and 120 hpf were incubated with acridine orange (AO), a fluorescent vital dye that labels dying cells. There was no significant difference between the number of AO-positive foci in wild-type and *hht* larvae at 24 hpf but AO-positive foci in *hht* larvae increased dramatically between 24-36 hpf (**Fig. 9A**). After peaking at 36 hpf, the number of detectable foci in *hht* continued to decline until 120 hpf. In contrast, the number of AO-positive foci in wild-type larvae remained consistently low and was significantly less than in *hht* larvae - at the 36 hpf peak, the number of AO-positive foci in *hht* was approximately 4-fold higher than that in wild-type. AO-positive cells in *hht* larvae accumulated predominantly in the most numerous cell types – those in the brain, eyes, and spinal cord (**Fig. 9B-C**)

**Figure 9.**
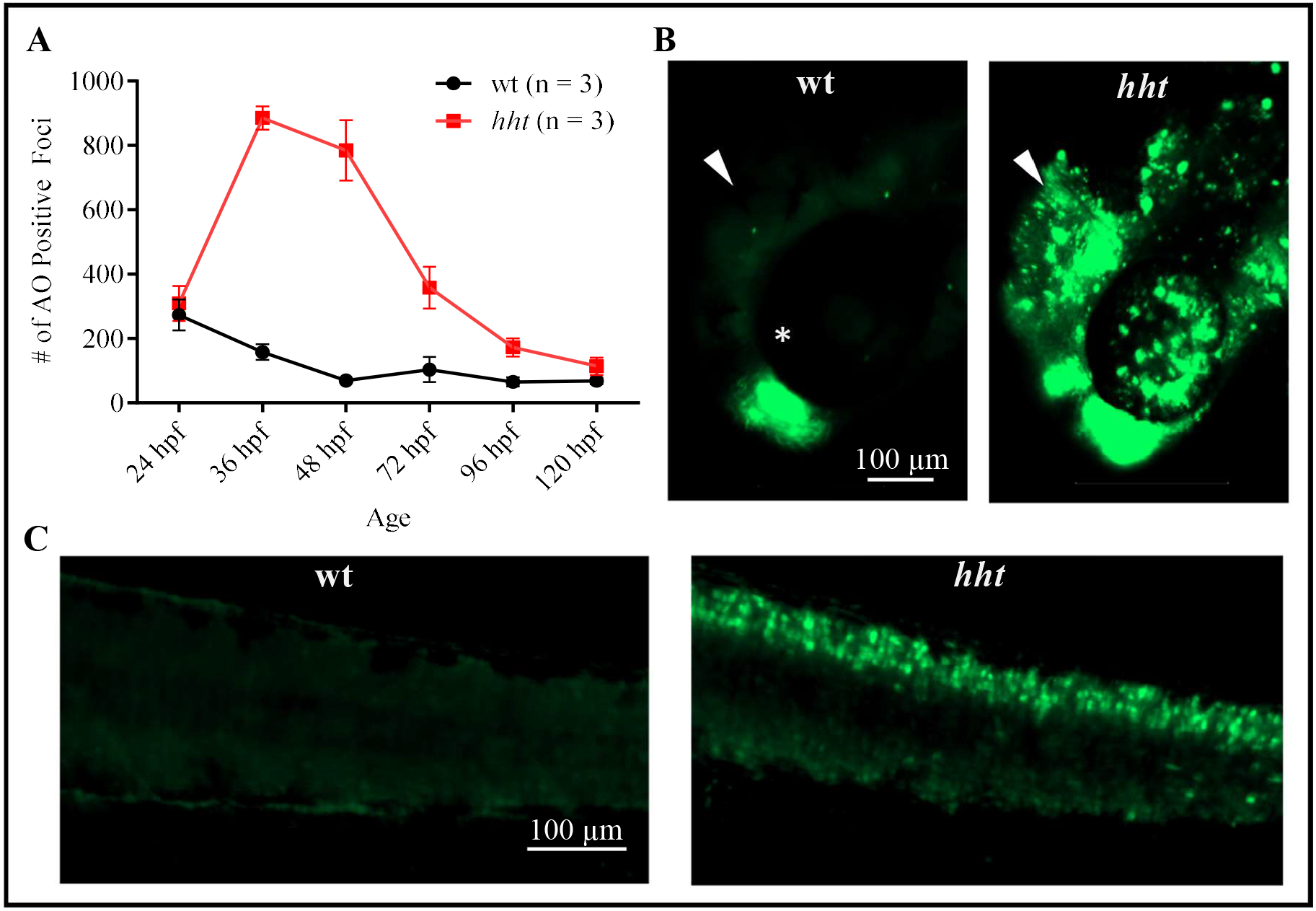
Acridine orange staining reveals increased cell death in the brain, eyes, and spinal cord of *hht* larvae. Cell death was assessed by counting the number of acridine orange-positive foci in 3 wild-type and 3 *hht* larvae.**A)** Cell death in *hht* was higher than in wild-type, peaking at 36 hpf. **B)** Cell death localized to the eyes and brain of 48 hpf *hht*. **C)** Cell death in the spinal cord in 48 hpf *hht*. Error bars = standard error of the mean. White arrow: brain, white asterisk: eye.

To assess the induction of double strand DNA breaks, wild-type and *hht* larvae at 24, 36, 48, 72, 96, and 120 hpf were probed with an antibody specific for γ-H2AX, the phosphorylated form of H2AX, a member of the H2A histone family. H2AX becomes phosphorylated on serine139 in response to double-stranded DNA breaks (DSB, Kuo and Yang 2008). At all ages examined, less than 30 positive foci were detected per wild-type larva. In contrast, the number of positive γ-H2AX foci in *hht* was indistinguishable from wild-type at 24 hpf, but increased significantly, from 8-to 13-fold, between 36-120 hpf. (**Fig. 10A**). γ-H2AX staining predominated in the brain, eyes, and spinal cord of *hht*, following the pattern of cell death as assessed by AO (**Fig. 10B-C**).

**Figure 10.**
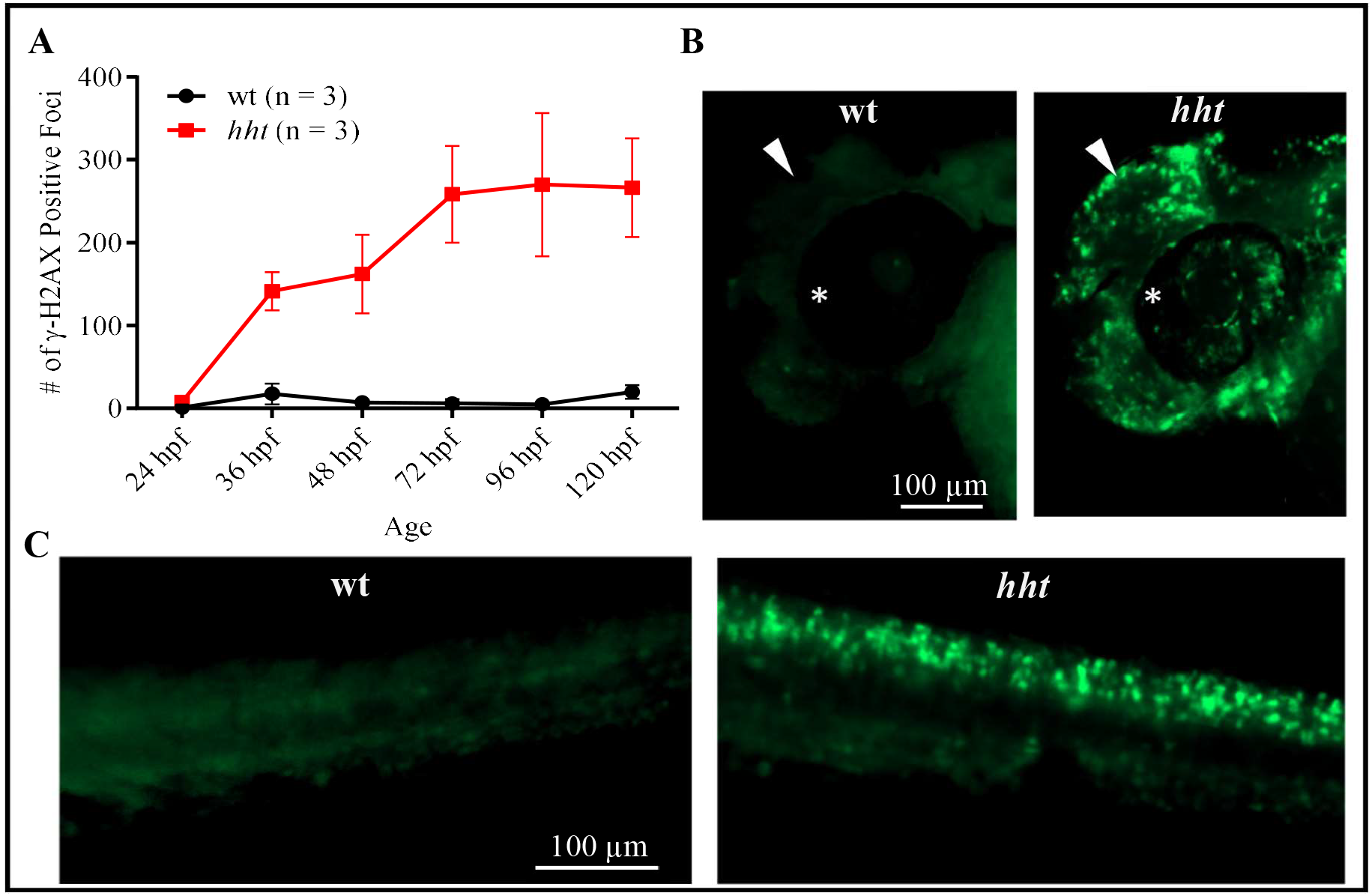
DNA damage is increased in *hht*. Double-stranded breaks were assessed by fluorescent antibody staining of γ-H2AX. **A)** DNA damage was minimal in wild-type, but striking in *hht* fish from 36-120 hpf. In 48 hpf *hht* fish, DNA damage was localized to the brain and eyes (B) and spinal cord (C). Error bars = standard error of the mean. White arrow: brain, white asterisk: eye.

### *Wild-type maternal* pola2 *mRNA sustains survival of* pola2*-deficient mutants*

To explain the extended survival of *pola2* null mutants of zebrafish compared with the *pola2*-deficient phenotypes of other model organisms, we hypothesized the presence of wild-type maternal *pola2* mRNA contributed from the wild-type chromosome of the heterozygous mother’s primary oocytes. Wild-type Pola2 protein originating from translation of wild-type maternal *pola2* mRNA would then support normal DNA replication during early embryogenesis and could be expected to be progressively diluted as cells divide in the absence of the wild-type gene in *hht* mutants. This scenario explains the relatively normal appearance of *hht* fish at 24 hpf, followed by the increasing disruption of development and eventual death at days 5-7 that characterize *hht* fish. The similarity in *hht* and chemically-inhibited fish phenotypes only after 24 hpf suggested are consistent with active DNA replication before 24 hpf in *hht* homozygotes.

To determine whether maternal wild-type *pola2* mRNA facilitates the survival of *hht* larvae, cDNA from 3 homozygous wild-type and 3 *hht* embryos at 2.5, 6, 24, 48, and 72 hpf were analyzed by quantitative allele-specific PCR to detect the presence of wild-type maternal *pola2* mRNA (**Fig. 11**). At 2.5 hpf, wild-type *pola2* transcripts were detected at similar levels in wild-type and *hht* embryos (*n* = 3; p > 0.5, t-test). By 6 hpf, the quantity of wild-type *pola2* transcripts in *hht* embryos was significantly reduced compared to wild-type embryos (*n* = 3; p < 0.05, t-test). Wild-type *pola2* mRNA was not detected in *hht* larvae at or beyond 24 hpf. Taken together, our results are consistent with a model in which the presence of wild-type *pola2* mRNA and protein is responsible for the sustained survival of the *hht* mutants, and that the degree of depletion of *pola2* mRNA and protein would be dependent upon the number of cell divisions in a given lineage.

**Figure 11.**
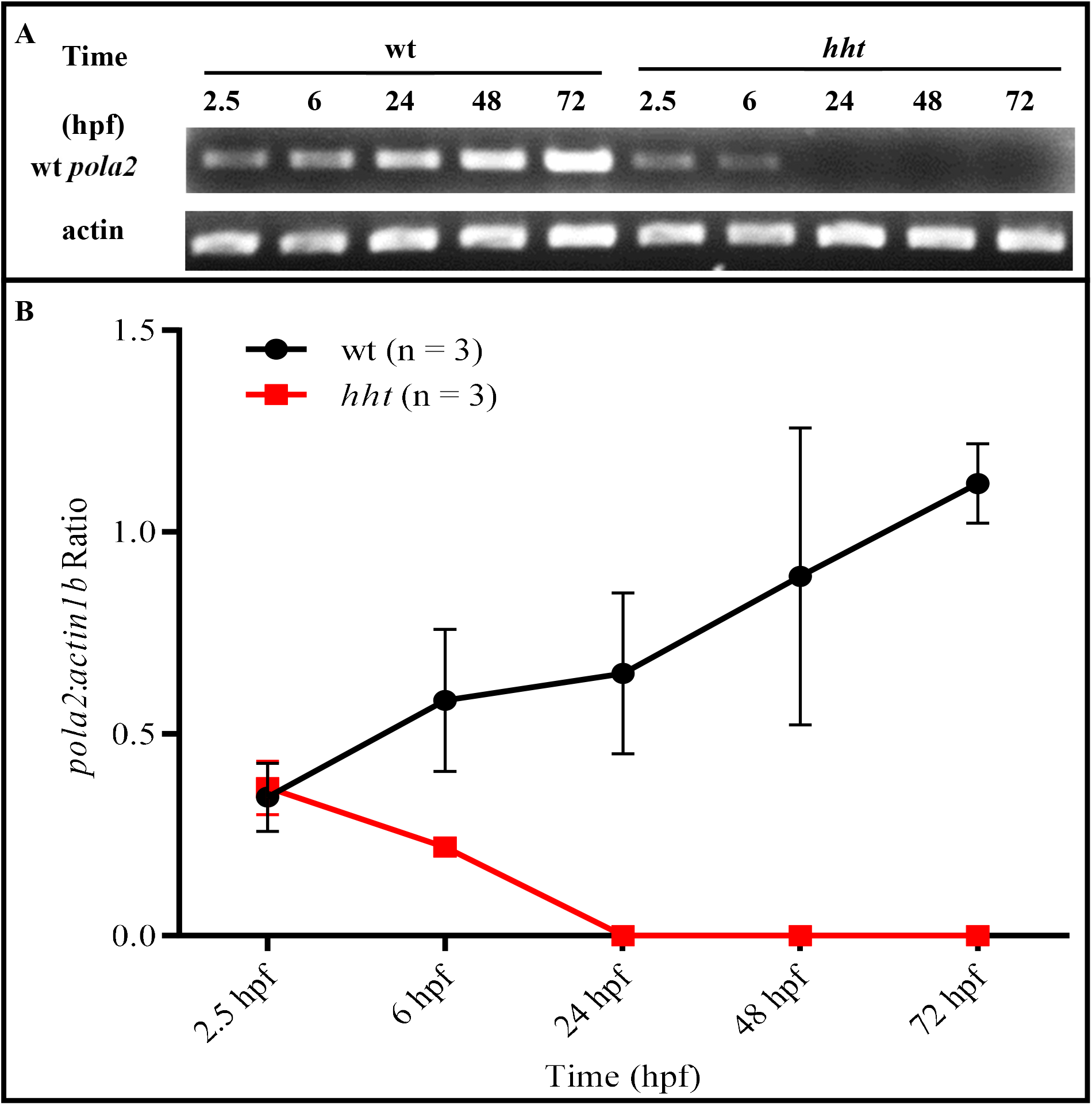
Wild-type *pola2* mRNA in *hht* embryos becomes undetectable by 24 hpf. **A)** Wild-type *pola2* transcripts were detected in *hht* by allele-specific primers at 2.5 and 6 hpf. Wild-type transcripts were not detectable in *hht* after 24 hpf. **B)** Wild-type *pola2* transcripts were normalized to *actin1b*. Wild-type *pola2* transcripts in *hht* embryos were present at comparable levels to wild-type embryos at 2.5 hpf but were significantly reduced compared to wild-type embryos by 6 hpf (p < 0.05)

## Discussion

Despite the clinical importance of nuclear atypia in the diagnosis and prognosis of human cancers, its mechanistic origins have been unclear. Hyperchromatic hematoxylin staining, often associated with increased prominence of nucleoli and irregular nuclear shapes, is common in nuclear atypia, consistent with cancer’s frequent association of aneuploid hyperploidy (Rosai 2004). The death and nuclear fragmentation of brain and retinal cells without detectable prior atypia is consistent with their normally small, dense nuclei, which makes heterogeneity in chromatin density inapparent. In contrast, the nuclei of larger cells such as those of the gastrointestinal tract and organs, are far larger. These larger cells contain the same amount of DNA per cell as cells with small nuclei, allowing variations in nuclear density to be more readily apparent. The atypical nuclei of *hht* gut epithelium are consistent with hyperploid aneuploidy, which is expected with DNA replication arrest in S phase. Cells trapped in S phase in *hht* mutant cells that are still alive would, by definition, contain more than 2n DNA content, and therefore be more darkly stained. Our genetic and chemical data, considered in the context of the association of the atypia associated with ionizing radiation and viral (in particular, papillomavirus) infection (Blum 1978, Stenbäck 1978, Carr and LiVolsi 1989, Kim *et al*. 2016, Sanfrancesco *et al*. 2013, Kufe *et al*. 2003), indicate that replicative stress may play the key role in nuclear atypia. Notably, each of these sources of replicative stress are associated with mutation (Adewoye *et al*. 2015, Gershenson 1986, Santos *et al*. 2011, Hanft *et al*. 2000, Zeeland *et al*. 1982), which, in turn, is necessary for the development of cancer (Tomlinson *et al*. 1996, Moolgavkar *et al*. 1981, Loeb *et al*. 1990).

The cell-dependent differences in phenotypes caused by *pola2* deficiency in *hht* mutants can be attributed to differences in replicative demand prior to the observed cell state. Since homozygous *hht* mutants cannot generate new wild-type *pola2* mRNA, the quantity of wild-type *pola2* mRNA, and presumably wild-type Pola2 protein, is diluted after each round of cell division (**Fig. 12**). As shown in our RT-PCR experiment, wild-type *pola2* mRNA is undetectable in *hht* fish by 24 hpf, leaving only the wild-type Pola2 protein produced before the maternal-to-zygotic transition, to support growth thereafter. Assuming the quantity of wild-type Pola2 protein in the progenitor cells for every tissue was the same, and is diluted with each cell division, tissues with a larger number of cells must have undergone more cell divisions and would therefore contain less wild-type protein per cell. The striking difference in the cellular disorganization of retinal cells and the organization of sensory epithelium in the *hht* mutant is readily explained by the retinal cells’ continuing replication vs. the sensory epithelium having reach terminally differentiation by about 24 hours (Dyballa *et al*. 2017). Phenocopying of a proliferation-dependent pattern of pleiotropy through the timed addition of chemical inhibitors of DNA synthesis is consistent with differential proliferation rates as the explanation for differential cellular responses to replicative deficiency.

**Figure 12.**
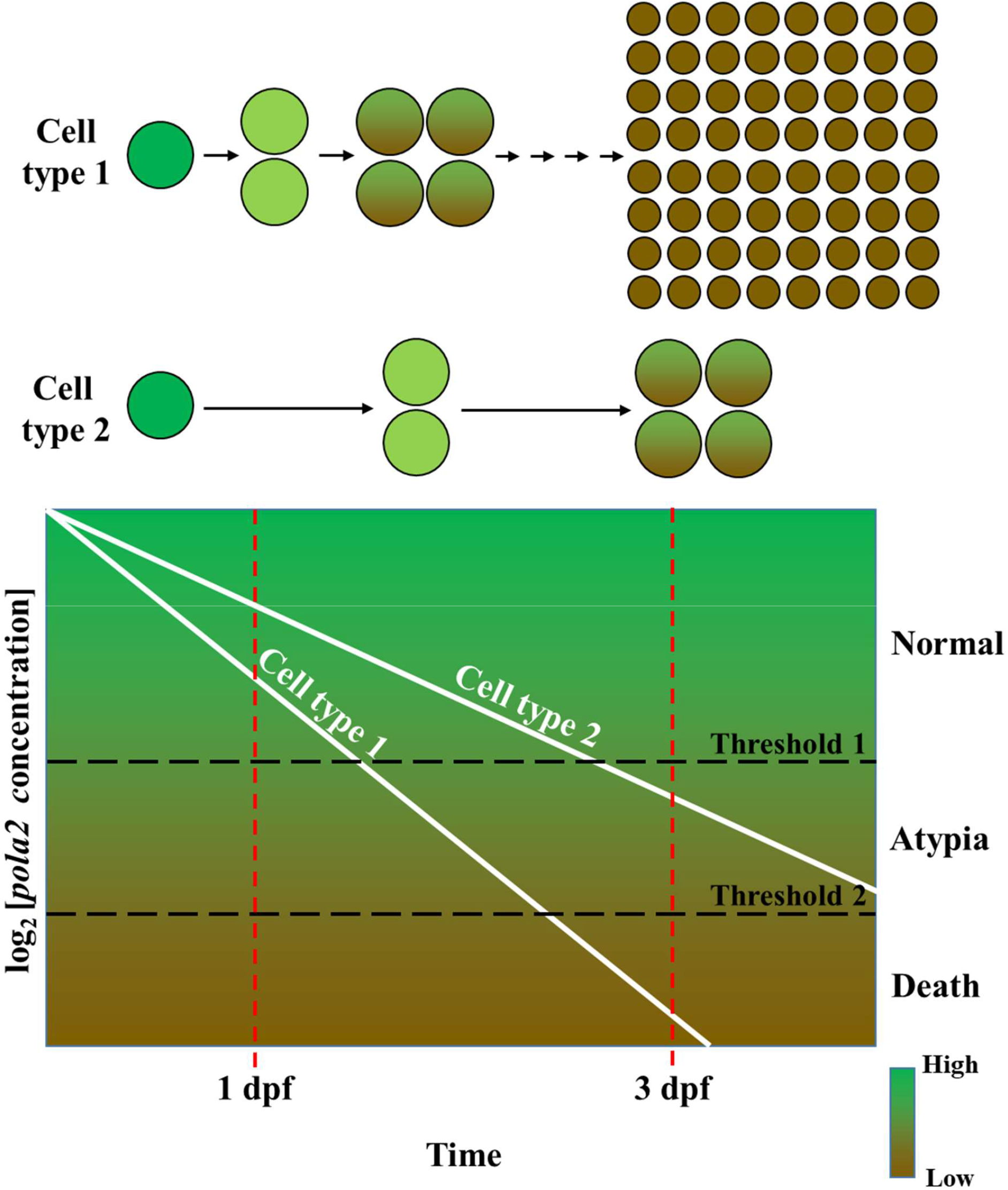
Differential dilution of *pola2* accounts for tissue-specific differences in nuclear atypia and cell death. Homozygous *hht* mutant embryos cannot generate wild-type *pola2* mRNA. The quantity of wild-type *pola2* mRNA and protein, is diluted after each round of cell division. In this model, highly proliferative cell types undergo more cell divisions per unit time, resulting in lower concentrations of wild-type *pola2* mRNA or protein per cell compared to cell types that proliferate slowly. Nuclear atypia is apparent in cells with large nuclei but inapparent in cells with small, condensed nuclei.

The *pola2* gene encodes the B subunit of Pol α and does not itself exhibit any known enzymatic activity. Molecular studies in mouse revealed that the B subunit is tightly associated with the zinc finger motifs in the carboxy-terminal domain of the catalytic subunit A (Mizuno *et al*. 1999). It functions as a “molecular tether” that recruits the catalytic polymerase subunit to the origin recognition complex (ORC) for the initiation of DNA replication (Collins *et al* 1993; Uchiyama and Wang 2004). Mutagenesis of the *POL12* gene, encoding the B subunit of Pol α in yeast, showed that mutations in the conserved C-terminus of the protein caused either lethality or temperature sensitivity while mutations in the non-conserved regions in the N-terminus had no effect (Foiani *et al*. 1995). The B subunit has been shown to be phosphorylated and dephosphorylated in a cell cycle-dependent manner and this phosphorylation is dependent on its association with the catalytic subunit (Foiani *et al* 1995; Ferrari *et al* 1996). The ability of Pol α to initiate DNA replication is maximized when the B subunit is phosphorylated while the N-terminus of the catalytic subunit A remains unphosphorylated (Schub *et al*. 2001). Other studies have shown that the B subunit pays a role in telomere maintenance in yeast (Grossi *et al*. 2004) and reprogramming the regenerative potential of the tail fin of adult zebrafish (Wang *et al*. 2019).

In non-vertebrate models, inactivating *pola2* mutations can result in immediate growth arrest (*S. cerevisiae*: Collins *et al*. 1993, Foiani *et al*. 1994, Foiani *et al*. 1995; *Arabidopsis*: Yang *et al*. 2009). Despite the high likelihood that the *hht* zebrafish are null mutants, they exhibit an extended lifespan of 5-7 dpf. As predicted by our replicative deficiency model, chemical inhibition of DNA replication in wild-type embryos starting at 1 dpf phenocopied *hht* at 3 dpf. Treatment during early embryogenesis caused death in wild-type larvae by 1 dpf, suggesting that normal DNA replication is indeed required during early embryogenesis for the survival of *hht* mutants, just as it is required in yeast and *Arabidopsis*. The detection of wild-type *pola2* mRNA in homozygous mutant embryos supports our hypothesis that DNA replication during embryonic stages of the mutants requires wild-type *pola2* mRNA. Since the homozygous mutant embryos have no genomic source for wild-type *pola2* transcripts, wild-type mRNA must have come from the heterozygous mother (fathers are also heterozygous but do not contribute mRNA to the oocyte). The maternal-to-zygotic transition (MZT) designates the time at which the zygotic genome is activated, and maternal transcripts become destabilized and rapidly degraded. In zebrafish, the transcription of the zygotic genome begins around 3 hpf, and most of the maternal transcripts are degraded by 6 hpf (Kane and Kimmel 1993, Schier 2007, Tadros and Lipshitz 2009). The time interval between MZT and onset of the *hht* mutant phenotype suggests that persistence of the wild-type protein is responsible for the survival of the mutant. That the pattern of subsequent somatic depletion of wild-type Pola2 protein in *hht* follows the pattern of DNA damage is consistent with our model for explaining the cell-specific pattern of *hht*’s pleiotropic mutant phenotype.

The extended lifespan of our *pola2* mutant, *hht*, and its development of time- and tissue-specific cellular abnormalities created an opportunity to explore the potential role of the progressive loss of DNA replication in the causation of an important cancer phenotype, nuclear atypia, and more generally, a frequent consequence of single gene mutations, pleiotropy. We showed that the progressive pattern of phenotypic change in *hht* can be explained by the initial presence of wild-type mRNA, followed by its progressive loss in genotypically homozygous mutant zygotes. In the course of our work, transcriptomic studies of wild-type, unfertilized zebrafish eggs showed that over 10,000 maternal transcripts are present that include essential genes such as *pola2* (Rauwerda *et al*. 2016). The present studies suggest that zebrafish mutants for maternally expressed essential genes may provide an opportunity to assess, without bias, both the tissue-specific and potentially pleiotropic effects of the progressive loss of gene function in the context of the whole animal. It is worth noting that the tissue-specific phenotypes of *hht* were discovered here using histology, but that whole-organism, 3-dimensional, pan-cellular, cell-resolution imaging method such as x-ray histotomography, will be better-suited for the quantitative study of volumetric phenotypes such as cell and tissue volume and shape across cell types and tissues (Ding *et al*. 2019). The incorporation of such approaches in the systematic study of pleiotropy may help us to understand patterns of cell and tissue-specific phenotypes that emerge in the course of disease.

## Materials and Methods

### Fish lines, mating, and embryo collection

Generation of ENU-mutagenized mutant *hht* was previously described (Mohideen *et al*. 2003). The background strain was a wild-type fish line acquired from the Connors fish farm, which differs from the Tu and AB laboratory strains that are in common use. The *hht* line is maintained as heterozygotes due to the larval-lethal nature of the mutation. Mating was carried out by placing male and female heterozygotes in Aquatic habitat tanks with dividers the afternoon prior to egg collection. Collected eggs were disinfected in 10% Ovadine (Syndel) for 1 min at room temperature then washed 3 times in charcoal-filtered water. Larvae were incubated at 28.5° C to maintain consistent speed of development.

### Histology

Procedure for zebrafish histology has been previously described (Copper *et al*. 2018) Larvae were dechorionated prior to submersion in 10% Neutral Buffered Formalin (Fisher) at 4° C. Samples were fixed overnight at room temperature with gentle agitation and subsequently embedded in agarose. For long-term storage, fixative was replaced with 70% ethanol. Agarose-embedded larvae are subsequently embedded in paraffin, sectioned, and mounted on glass slides. Sections were stained with hematoxylin and eosin and imaged with Zeiss Axio Imager M.2.

### X-ray histotomography

3-dimensional images by micro-CT were extracted from the 3d.fish database created by Ding *et al*. Links to the exact images presented in Figure 2 are provided below.

3 dpf wt: http://3d.fish?s=wt_3dpf&r=0&z=sagittal&c=0.09086888387667061,0.07090368892198975,0.4822530864197531,0.23534955311213995,0&i=518,317,671&bri=100&con=100&ol=

5 dpf wt: http://3d.fish?s=wt_5dpf&r=0&z=sagittal&c=0.026697595375619665,0.0606159072099105,0.3721088629782046,0.18159687740134256,0&i=152,416,742&bri=100&con=100&ol=

3 dpf *hht*: http://3d.fish?s=hht_3dpf&r=2&z=sagittal&c=0.08014927360645446,0.3082981561232899,0.8196371398078974,0.39999999999999997,0&i=647,485,430&bri=100&con=100&ol=

5 dpf *hht*: http://3d.fish?s=hht_5dpf&r=0&z=sagittal&c=0.02394390397352475,0.17317734966732362,0.44653063557384554,0.2179162528816111,0&i=210,471,293&bri=100&con=100&ol=

### Sequencing

mRNA was extracted from 72 hpf wildtype and *hht* larvae using the RNeasy Mini Kit (Qiagen) and reverse transcribed to cDNA. PCR was performed on cDNA with 3 sets of primers:

-23∼596:

Forward (POLA2F1: TTGAACATCAGAGGACAATA)

Reverse (POLA2R1: TCTCCTCCGTCCAGCATCTC)

503∼1274:

Forward (POLA2F2: AGAGGTGGTTTCCACATTTG)

Reverse (POLA2R2: ACAATACCAACTTGCATGC)

1180∼1823:

Forward (POLA2F3: ACCAGGTGACAGAAACATTT)

Reverse (POLA2R3: TAAAGTTCAAACATTGTATG).

PCR was performed on gDNA with 1 set of primer for exon 2 of *pola2*:

Forward (POLA2F1: TTGAACATCAGAGGACAATA) anneals to the 5’ end of the 2^nd^ exon of *pola2*.

Reverse (POLA2gR: TGACTCCAAACAATGTTGTACTTTGATAGTCATTTG) anneals to the 2^nd^ intron of *pola2*.

PCR products were purified with the QIAquick PCR purification kit (Qiagen) and submitted to Genewiz for sequencing.

### Genotyping

Allele-specific primers were designed based on the SNAP primer method described by Drenkard *et al*. Forward primers span the AC repeats. Wild-type specific primers detect alleles with only two repeats and mutant specific primers detect alleles with three repeats.

Wild-type forward primer (POLA2wtF: CACTTTCAACATAGCCTTCAACAATGACAGC); Mutant forward primer (POLA2mtF: GACACTTTCAACATAGCCTTCAACAATGAGACA); gDNA reverse primer

(gPOLA2ASR: TGACTCCAAACAATGTTGTACTTTGATAGTCATTTG). cDNA Reverse primer: (cPOLA2ASR: GCTCCAACATTTTGTCGATGGTGCC)

### RNA rescue

Wild-type and mutant *pola2* mRNA was extracted from homozygous wild-type or homozygous mutant embryos, respectively, from a 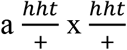 cross with the RNeasy Mini Kit (Qiagen) and reverse transcribed into cDNAs using the M-MLV Reverse Transcriptase kit (ThermoFisher). cDNA from wild-type embryos were subjected to allele-specific PCR to eliminate heterozygotes. cDNAs were amplified by PCR with the following primers:

Forward primer (POLA2-BglIIF): <u>AGATCT</u>TTGAACATCAGAGGACAATA

Reverse Primer (POLA20MluIR): <u>ACGCGT</u>TAAAGTTCAAACATTGTATG

and cloned into PCR®II-TOPO® vectors for sequencing. After sequences are verified, the inserts are subcloned into PT3TS(4) vectors. Plasmid DNA was extracted from competent cells with the QIAprep Spin Miniprep Kit (Qiagen) and linearized with Xmal. mRNA was generated from the linearized plasmids with the T3 RNA polymerase (NEB). 200 pg of wild-type or mutant *pola2* mRNA were injected into embryos from a 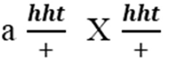 cross at the 1-cell stage. All injected embryos in the first round of rescue experiments were genotyped to ensure that no phenotypic wild-type larvae were genotypically *hht*.

### CRISPR/Cas9 knockout

Gene-specific primer was designed for *pola2* and annealed to a universal primer containing the sgRNA scaffold as described in protocol provided by Dr. Wenbiao Chen’s lab at Vanderbilt University. 1 μL of 10 μM gene-specific primer was mixed with 1 μL of 10 μM sgRNA scaffold primer, 2.5 μL 2X NEB buffer 2 with BSA and annealed at 98° C for 1 min then cooled to 37° C with a ramp speed of 0.1° C/sec. 0.5 μL of 500 μM dNTP and 0.5 μL of T4 DNA polymerase were added to the mixture and incubated for 20 min. T4 DNA polymerase was then inactivated by incubation at 75° C for 20 min. sgRNA was synthesized using the MaxiScrip T7 kit (ThermoFisher Scientific). 1 μL of Turbo Dnase I was added and incubated at 37° C for 15 min to remove residual DNA primers. sgRNA was purified using the mirVANA kit (ThermoFisher Scientific). 600 ng/μL Cas9 protein was mixed with either 100 ng/μL *pola2* sgRNA and injected into wild-type zebrafish embryo at 1-cell stage.

pola2KO: ATTAATACGACTCACTATAGGGGTCCGCGCTGGATGGGGAAgttttagagctagaaatagc; sgRNA scaffold: ttttgcaccgactcggtgccactttttcaagtTgataaCggactagccttattttaacttgctatttctagctctaaaac

### Cell cycle analysis

20 wild-type and *hht* larvae were collected at 3 and 5 dpf and dissociated into single cells. Cells were resuspended in 2 mL 70% EtOH at room temperature for 10 min to fix then stored at 4° C before use. Cells were centrifuged at 500 x g for 5 min to remove supernatant. 1 mL propidium iodide solution from DNA QC Particles Kit (BD) was added to each cell mixture. Cells were sorted with FACSCalibur (BD) and cell cycle was analyzed using ModFit LT V3.3.11.

### EdU staining

For labeling of proliferating cells, 2, 3, 4, and 5 dpf wildtype and *hht* larvae were incubated for 30 min at 28.5° C in 400 µM EdU solution (Life Technologies). Larvae were incubated in charcoal-filtered water for 30 min at 28.5° C, then fixed immediately in cold 10% neutral buffered formalin. Detection of EdU incorporation was performed according to manufacturer’s instructions. Samples were visualized with Axio Zoom.V16 fluorescence stereo microscope (Zeiss).

### Chemical inhibition

Wild-type embryos were submerged in charcoal-filtered water, hydroxyurea (Sigma-Aldrich), or aphidicolin (Sigma-Aldrich) at 0, 2, 6 hpf or 24 hpf. For the 24 hpf treatment, fresh solutions were administered at 48 hpf. Hydroxyurea was dissolved in deionized water to prepare a 500 mM stock and aphidicolin was dissolved in DMSO to prepare a 10mM stock. Stock solutions were diluted to appropriate concentrations for each experimental condition so that equal volume was added.

### Acridine orange staining

24, 36, 48, 72, 96, and 120 hpf wildtype and *hht* larvae were incubated in 2 μg/mL acridine orange for 30 min at 28.5° C, then washed for 5 min in charcoal-filtered water for 5 min at 28.5° C. Larvae were visualized with Axio Zoom.V16 fluorescence stereo microscope (Zeiss).

### γ-H2AX staining

24, 36, 48, 72, 96, and 120 hpf wildtype and *hht* larvae were fixed in cold 10% neutral buffered formalin. Samples were permeablized in acetone at -20 °C for 7 min then washed with distilled water for 5 min followed by two washes in PBS for 10 min. Samples were then blocked in 5% goat serum for 1 hr at room temperature and incubated in 1:100 γ-H2AX primary antibody (Genetex) overnight at 4 °C. After 3 washes in PBST for 15 min, samples were incubated in 1:1000 Alexa Fluor 488 goat anti-rabbit secondary antibody (Life Technologies) for 3 hrs at room temperature. After 3 washes in PBST for 15 min, samples were visualized with Axio Zoom.V16 fluorescence stereo microscope (Zeiss).

### Wildtype *pola2* transcript detection

DNA and RNA were extracted from 2.5, 6, 24, 48, and 72 hpf embryos with AllPrep DNA/RNA Mini Kit (Qiagen). Embryos were genotyped with allele-specific PCR. RNA of genotyped homozygous wild-type and *hht* embryos were reverse transcribed with oligo-dT. Quantitative PCR was performed on cDNA with allele-specific primers for wild-type *pola2* and *actinb1*:

Forward (Actinb1F: CATCCGTAAGGACCTGTATGCCAAC)

Reverse (Actinb1R: AGGTTGGTCGTTCGTTTGAATCTC)

as loading control.

PCR products from three homozygous wild-type and mutant embryos were subjected to electrophoresis. Relative quantity was calculated by densitometry as a ratio of *pola2* product:*actinb1* product.

## General

We thank Margaret Hubley, Gail Broda, and Kathryn Early for help in maintaining wild-type and mutant zebrafish lines and generating embryos; Jean Copper and Lynn Budgeon for help in development of zebrafish histology.

## Author contributions

The project was conceived by KCC; research design was by AYL and KCC; experiments were done by AYL and GKT; analysis was by AYL, GKT, and KCC; AYL and KCC wrote the paper; KCA, DBVR, and VAC contributed to experimental design and editing. Finally, we would like to thank Dr. Larry Loeb for his encouragement during the conception of this project.

## Supplementary Materials

**Supplemental S1.**
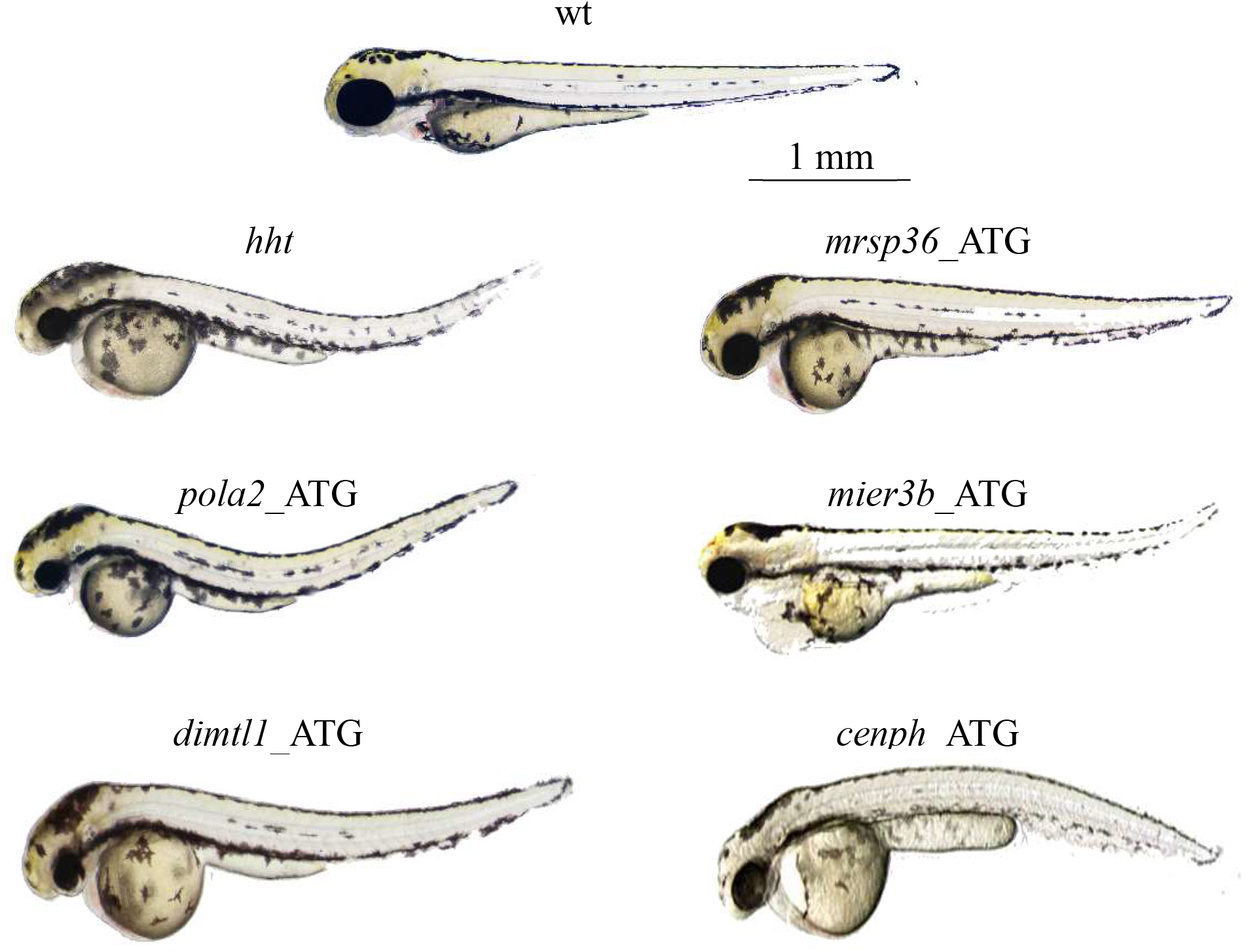
Morphants of the five candidate genes in the region identified by positional cloning, *mier3b, mrps36, cenph, pola2*, and *dimtl1*. Only *pola2* and *dimt1l* morphants exhibit a combination of small eyes, enlarged yolk, and dorsal curvature characteristic of *hht* mutants.

**Supplemental S2.**
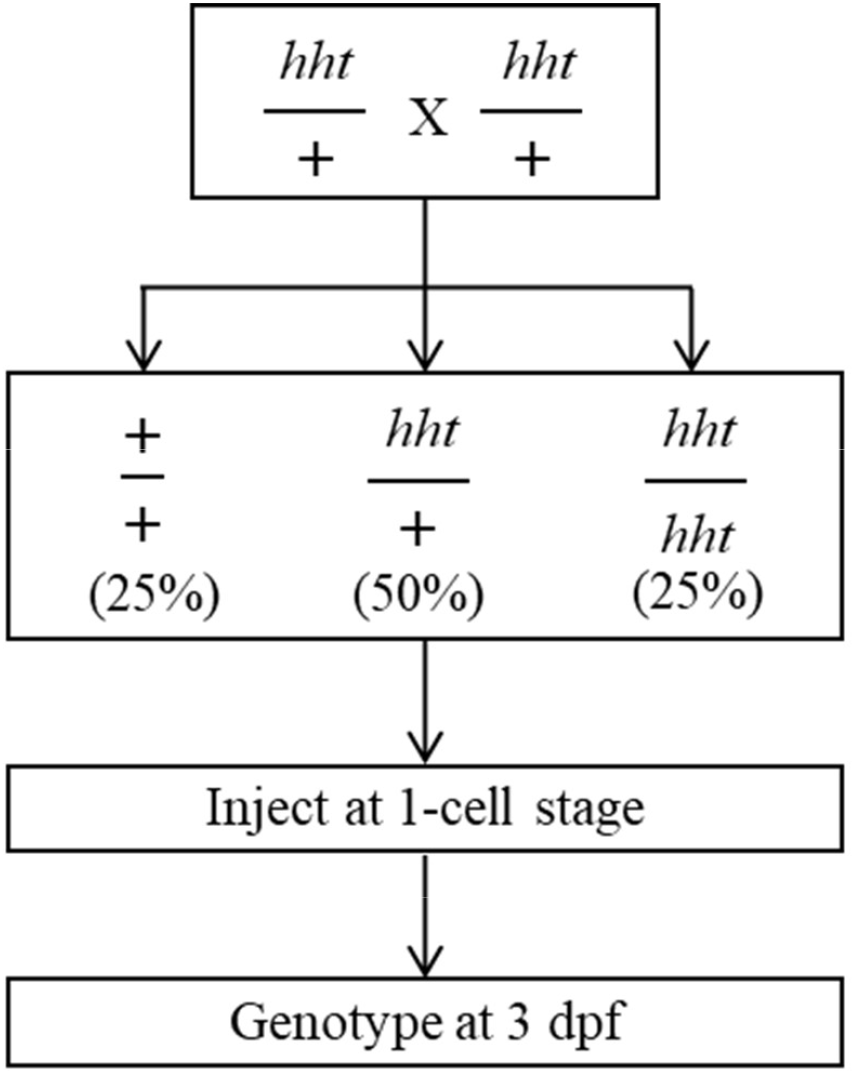
Wild-type *pola2* mRNA rescue schematic. Wild-type *pola2* mRNA was injected into 1-cell stage embryos from a 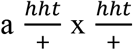 cross. Larvae were genotyped to confirm that rescued larvae were homozygous for the *hht* mutation.

**Supplemental S3.**
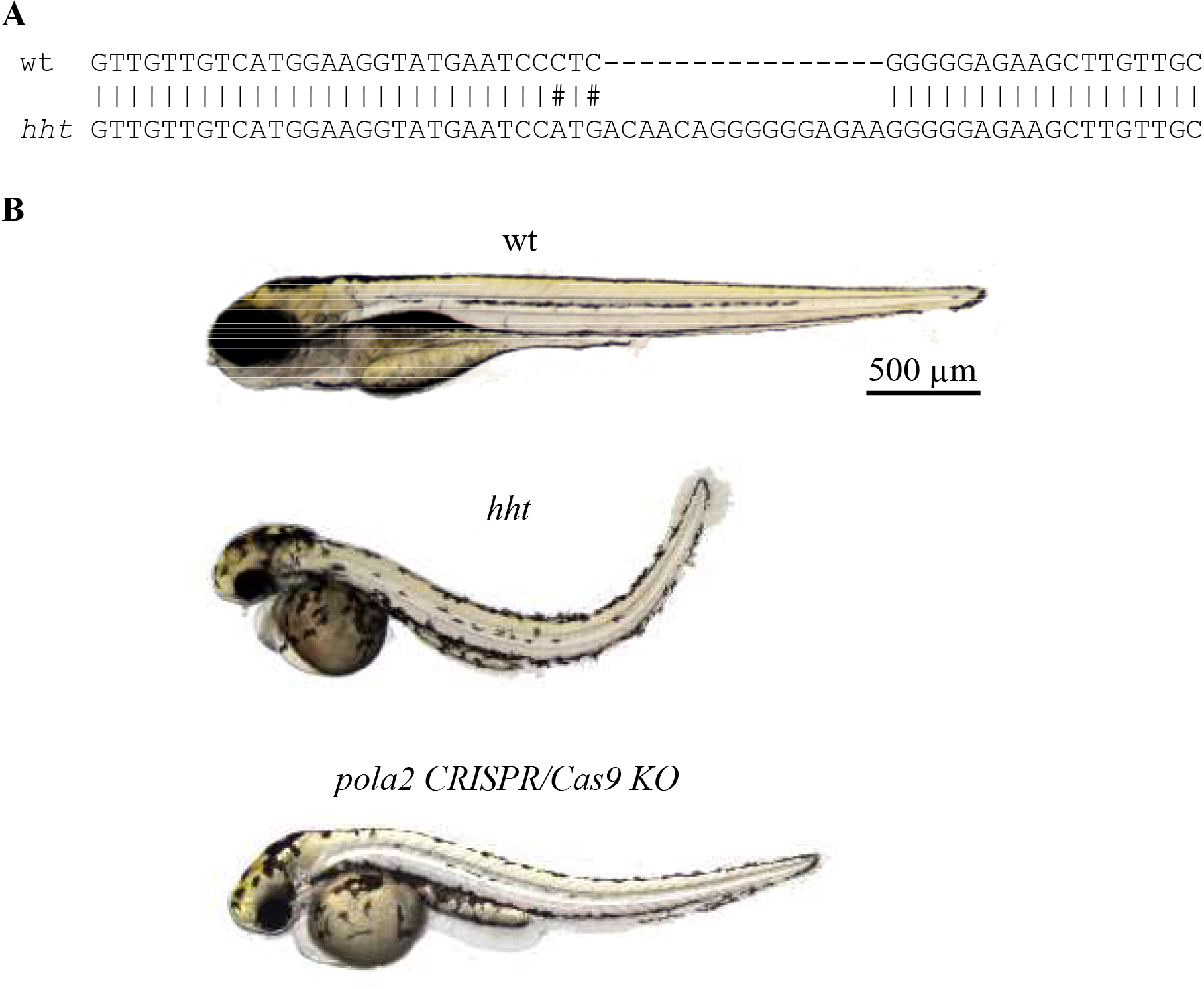
Shared phenotype of CRISPR/Cas9 *pola2* knockouts and *hht*. **A**) The CRISPR/Cas9 knockout carried a 16-nucleotide insertion between nucleotides 927-928, causing a frameshift that changes the protein sequence after amino acid 308 in the 600-amino acid zebrafish Pola2 protein. Differences in severity in phenotype are consistent with variation in genetic background. **B**) Representative images of wild-type, *hht*, and *pola2* CRISPR/Cas9 knockout zebrafish larvae at 5 dpf. *hht* and *pola2* knockout fish shared small eyes, small head, enlarged and rounded yolk, and dorsally curved body.

**Supplemental S4.**
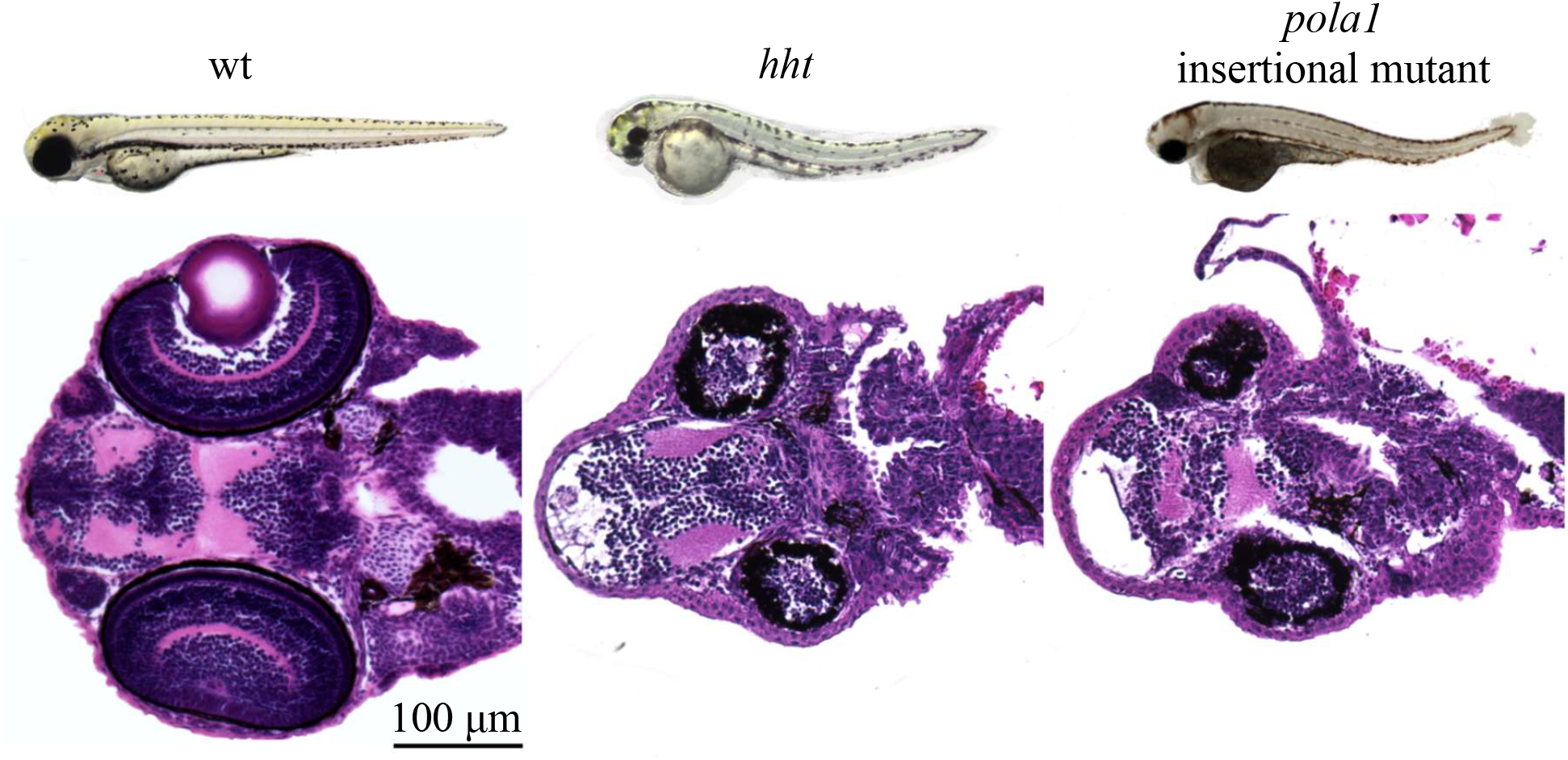
Mutants of *pola1* and *pola2* exhibit virtually indistinguishable gross and histological phenotypes. At 3 dpf, *hht*, our *pola2* null mutant, and a *pola2* insertional mutant both exhibit the combinatorial gross phenotype of small eyes, small head, rounded yolk, and curved body. Histologically, both mutants show a drastic reduction of cell number and loss of retinal layers in the eyes, reduced volume and disorganization of the white and gray matter of the brain. Scale bar = 100 μm.

**Supplemental S5.**
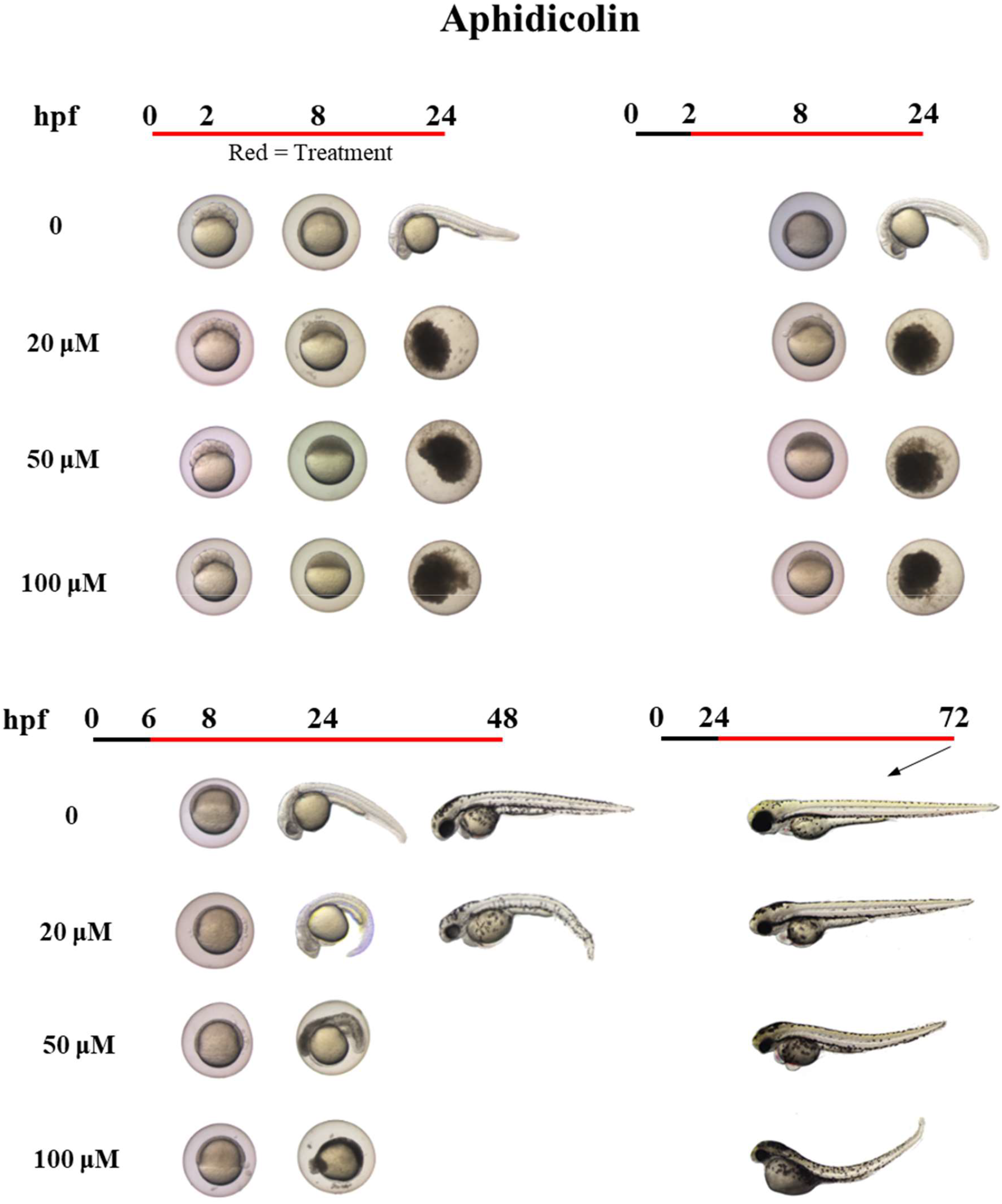
Inhibition of replicative DNA polymerases by aphidicolin beginning in wild-type zebrafish larvae at 24 hours, but not earlier, phenocopies *hht* at 72 hpf. Wild-type larvae were continuously exposed to 0, 20, 50, or 100 μM aphidicolin in DMSO. Treatment began at 0, 2, 6, or 24 hpf. 10 larvae were treated for each condition. Representative images are shown. Black line = no treatment; red line = treatment.

**Supplemental S6.**
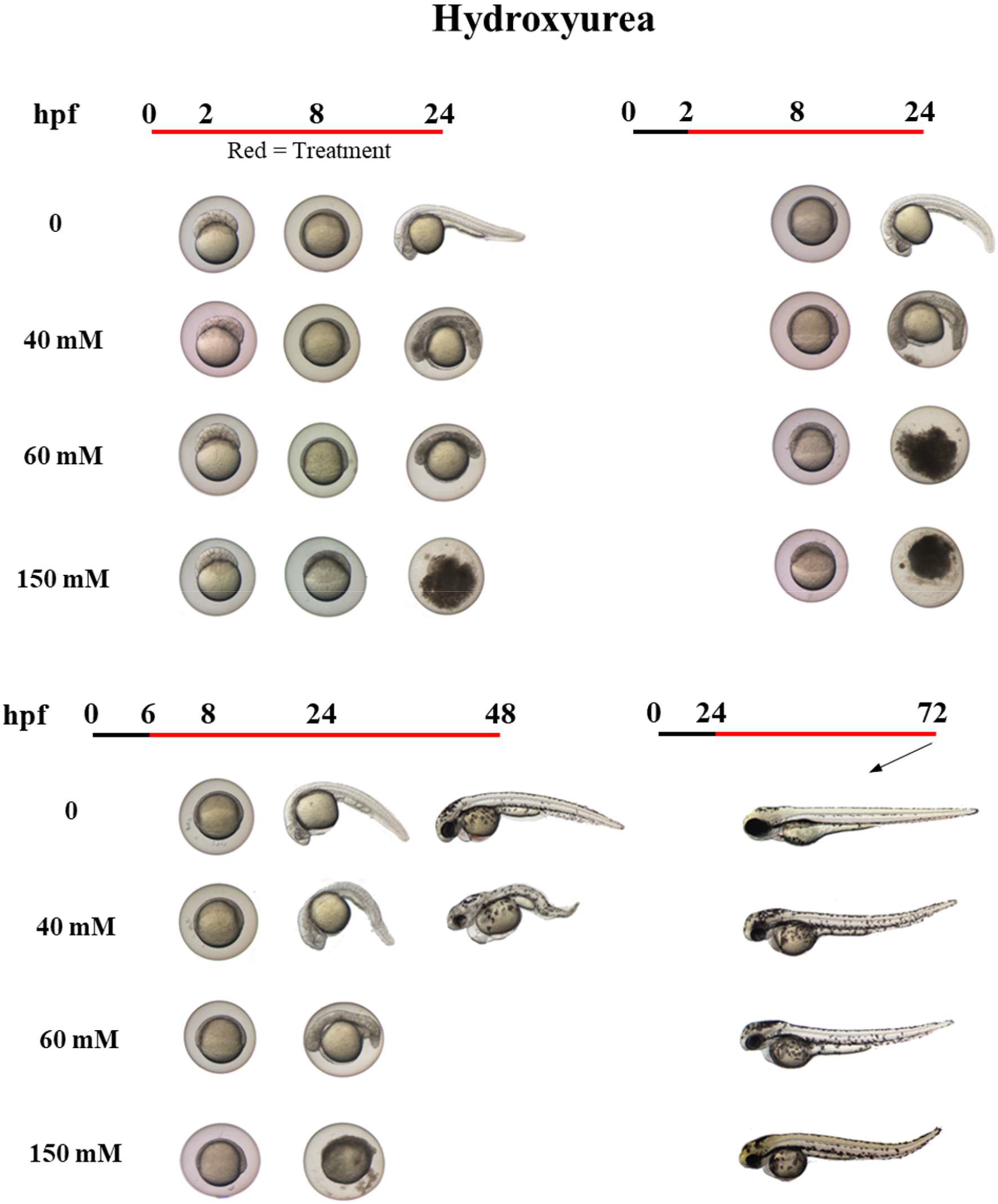
Inhibition of deoxyribonucleotide production by hydroxyurea in wild-type zebrafish larvae beginning at 24 hours, but not earlier, phenocopies *hht* at 72 hpf. Wild-type larvae were continuously exposed to 0, 40, 60, or 150 mM aqueous hydroxyurea. Chemical treatment began at 0, 2, 6, or 24 hpf. 10 larvae were treated for each condition. Representative images are shown. Black line = no treatment; red line = treatment.

